# Beyond the dyad: higher-order structure within cohesive animal groups reveals the social pathway to leadership

**DOI:** 10.1101/2022.05.30.494018

**Authors:** Lorenzo Betti, Federico Musciotto, Danai Papageorgiou, Federico Battiston, Damien R. Farine

## Abstract

Revealing the consequences of social structure in animal societies is largely determined by our ability to accurately estimate functionally relevant patterns of social contact among individuals. Of particular relevance are social contacts that drive and maintain the cohesion of moving animal groups. To date, studies have predominantly built up social structure from dyadic connections, which while effective in open societies—where groups are ephemeral—may not be sufficient to characterise the fine-grained structure within more cohesive animal groups. This is because associations or interactions can involve more than two individuals participating together, which current approaches cannot distinguish from independent sets of dyadic connections. Here we apply higher-order temporal network approaches to high-resolution GPS data from every group member in two cohesive groups of vulturine guineafowl (*Acryllium vulturinum*). By quantifying moment-by-moment association dynamics, we reveal clear sex-biased contributions to the group cohesion. Specifically, males form the cohesive core of the group as they participate in temporally consistent sub-groups and tend to occupy forward positions when in movement. Females instead remain peripheral and are more likely to leave sub-groups, while also occupying rear positions in moving sub-groups. These results reveal how the cohesion among males, which were previously found to be more likely to lead group movements, allows them to more often form a majority, confirming that self-organisation within the group can drive leadership patterns independently of dominance. Our study thus demonstrates that sub-group association dynamics derived from high-resolution GPS data can provide insights in how the fine-scale spatial and social organisation of cohesive groups, and their temporal dynamics, can explain key processes like leadership.

## INTRODUCTION

It is well-established that the repeated patterns of social contact among individuals can have far reaching consequences (Cantor et al. 2021). For example, how distributed social contacts are and how frequently they are formed shapes the spread of information (Voelkl & Noë 2008; Aplin et al. 2012) and disease dynamics (Silk et al. 2019; Ashby & Farine 2022). Further, the quality of relationships that individuals maintain can have consequences for their survival and that of their offspring (Alberts 2019; Gerber et al. 2022), while who they interact with can impact their ability to reproduce (Farine & Sheldon 2015; Fisher et al. 2016). Even though the consequences of the structure that arises from social contacts are evident, the field of animal social networks has only recently started looking beyond pairwise (or dyadic) and static patterns of social contacts, with greater focus on capturing aspects such as the temporality of networks as well as their higher-order properties (Silk, 2023). While the tools we have been using so far allow us to capture a meaningful approximation of underlying network structure for processes that are naturally dyadic, like disease transmission (Krause et al. 2013; Farine & Whitehead 2015), these tools may not capture some fundamental elements of the social glue that holds animal groups together. This is because human and many non-human animal societies are made up of a rich diversity of relationships, many of which are best represented by social units (e.g. a family unit, a society, and many levels in between; Camerlenghi et al. 2022; Grueter et al. 2020; Papageorgiou & Farine 2021; Camerlenghi & Papageorgiou 2025). Thus, our ability to capture the presence and functional importance of such social units—such as the substructure of animal groups with stable group membership—and how they contribute to group processes are limited. Here, we demonstrate the use of a group-level, temporal approach for extracting key social components from high-resolution tracking data.

Social networks—where nodes represent individuals and edges reflect the inferred relationship between the individuals—are widely used to describe the emergent properties of social groups or broader populations (Farine & Whitehead 2015; Sosa et al. 2021). In practice, constructing networks starts with observations of dyadic associations (two individuals present together) or dyadic interactions (either affiliative or agonistic). For example, individual giraffes have been defined as belonging to the same observed group when they forage or move together, but not when they move past each other in opposite directions, with groups defined as clusters of Masai giraffes (*Giraffa camelopardalis*) that are more than 500 m from other giraffes (Bond et al. 2021). In other examples, great tits (*Parus major*) (Psorakis et al. 2012; Aplin et al. 2015) and juvenile hihis (*Notiomystis cincta*) (Franks et al. 2020) are assumed to belong to the same group if observed visiting feeders together within the same feeding ‘wave’, and when constructing association networks, members of olive baboon (*Papio anubis*) troops are assumed to be interacting when they groom each other (Tung et al. 2015). While each of these methods differ in their biological definition, they generally follow the same principle of summarising observations of individual associations or interactions into rates (Whitehead 2008; Hoppitt & Farine 2018) forming the basis for generating social networks.

Social networks show their limitations when describing the social structure within cohesive social units. In species that live in stable groups that move cohesively, individuals spend most of their lives in very close proximity to each other. Within these groups (Battiston et al. 2020), genuine social preferences might be present among subsets of individuals, but these can be therefore masked because individuals come into frequent contact with all members of their group—both preferred and not—making it difficult to capture the social structure of the group. Yet fine-scale preferences may be important because social preferences among individuals can shape how groups make decisions. For example, Chacma baboons (*Papio ursinus*) have been shown to be more likely to follow the initiation attempts of their closest associates (King et al. 2011). In many animal groups, collective movement decisions are made by selecting the direction with the majority of initiators (see references in Farine et al. 2025), known as a ‘majority rule’ (Conradt & Roper 2003). Yet this majority rule is inherently a process involving clusters of close associates rather than sets of dyadic relationships.

A challenge for scaling up from dyadic to higher-order clusters of relationships is that it requires a massive increase in the amount of data necessary. Recent technological advances in GPS tags (He et al. 2022), such as automated reverse GPS-systems (Nathan et al. 2022), can now provide continuous, high-resolution positions for a large fraction of group members, which have already proven effective in understanding the social organization of animal groups. For example, data from a group of baboons where the majority of members were fitted with GPS collars was used to infer how groups make movement decisions (Strandburg-Peshkin et al. 2015) and how social relationships impact where individuals move next (Farine et al. 2016). In captivity, a high resolution tracking system applied to zebra finch (*Taeniopygia castanotis*) colonies was used to infer the establishment of breeding pairs over time (Maldonado-Chaparro et al. 2021), how the developmental environment affects the emergent structure of populations (Wang et al. 2022), and to identify the multitiered social environment of individuals (Zhang et al. 2025). By leveraging such coverage, high-resolution GPS can resolve the moment-by-moment formation and dissolution of associations among subsets of individuals within cohesive groups, providing a lens to uncover their internal social structure and processes. For example, in olive baboons the outcome of collective decisions were found to be best predicted by the relative sizes of clusters of initiators, and the study obtained these by aggregating dyadic leader-follower data into temporal, higher-order clusters (Strandburg-Peshkin et al. 2015). As these types of data become more widely collected, we will also benefit from advancing our use of analytical methods that can better capture the multidimensionality of the social relationships within social groups.

In this paper, we use continuous, high-resolution GPS tracking of two entire groups of wild vulturine guineafowl (*Acryllium vulturinum*) to quantify the dynamics of their association patterns and reveal the internal social structure of the group. Vulturine guineafowl are large and predominantly terrestrial birds that are endemic to East Africa. They are also highly social, living in groups of 11 to 65 individuals that move cohesively together (Papageorgiou et al. 2019). GPS-based tracking has revealed that these groups associate preferentially with specific other groups to form a multilevel society (Papageorgiou et al. 2019). However, while the population-level structure—the stability of group membership over time and the association among groups that form the upper two levels of this three-level society—is clearly evident (Farine 2025), capturing what happens within groups has been proven to be more challenging. To date, we know that groups contain multiple reproductive males and females (Papageorgiou et al. 2019), and that the society is male-structured, with all males being on the top of the dominance hierarchy (Papageorgiou & Farine 2020; Dehnen et al. 2022) and females being the dispersing sex (Klarevas-Irby et al. 2021). Groups make shared collective decisions using a majority rule, but while all group members can initiate movements, males (the philopatric sex) are disproportionately more likely to be in the largest cluster and, thus, be successful in leading a group movement (Papageorgiou et al. 2024). However some exceptions exist, as a distinct shift in the role of females (the dispersing sex) have been observed within the group. Notably, females receive less aggression from males after they breed successfully, which is likely because they become mothers to male group members (Dehnen et al. 2025).

The high degree of cohesion within vulturine guineafowl groups makes it challenging to uncover how social preferences are directed among members of the same group. This challenge also limits our ability to explain why males are more likely to lead because there are no effects of dominance within males (or females) (Papageorgiou et al. 2024). What we do often observe visually is that males move in extremely tight clusters (Fig. 1), which could make them more likely to end up in a majority cluster, suggesting that these within-group movement dynamics could play a central role to group cohesion. To investigate how such movement patterns contribute to group cohesion, we use the continuous 1 Hz GPS data on the movements and relative position of all the group members in two groups over 6 days. We then describe the composition and temporal stability of sub-groups, including quantifying how likely birds are to transition between sub-groups of different sizes. We also quantify spatial positions of birds within their sub-group. Because females disperse while males are philopatric and remain in their natal group (Klarevas-Irby et al. 2021; Klarevas-Irby & Farine 2024), we expect stronger cohesiveness among males as a result of the likely high within-male genetic relatedness, which may reinforce associations among males as a sub-group. We use this expectation as our a priori knowledge (Ferreira et al. 2020) to evaluate the performance of our approach.

**Figure 1.**
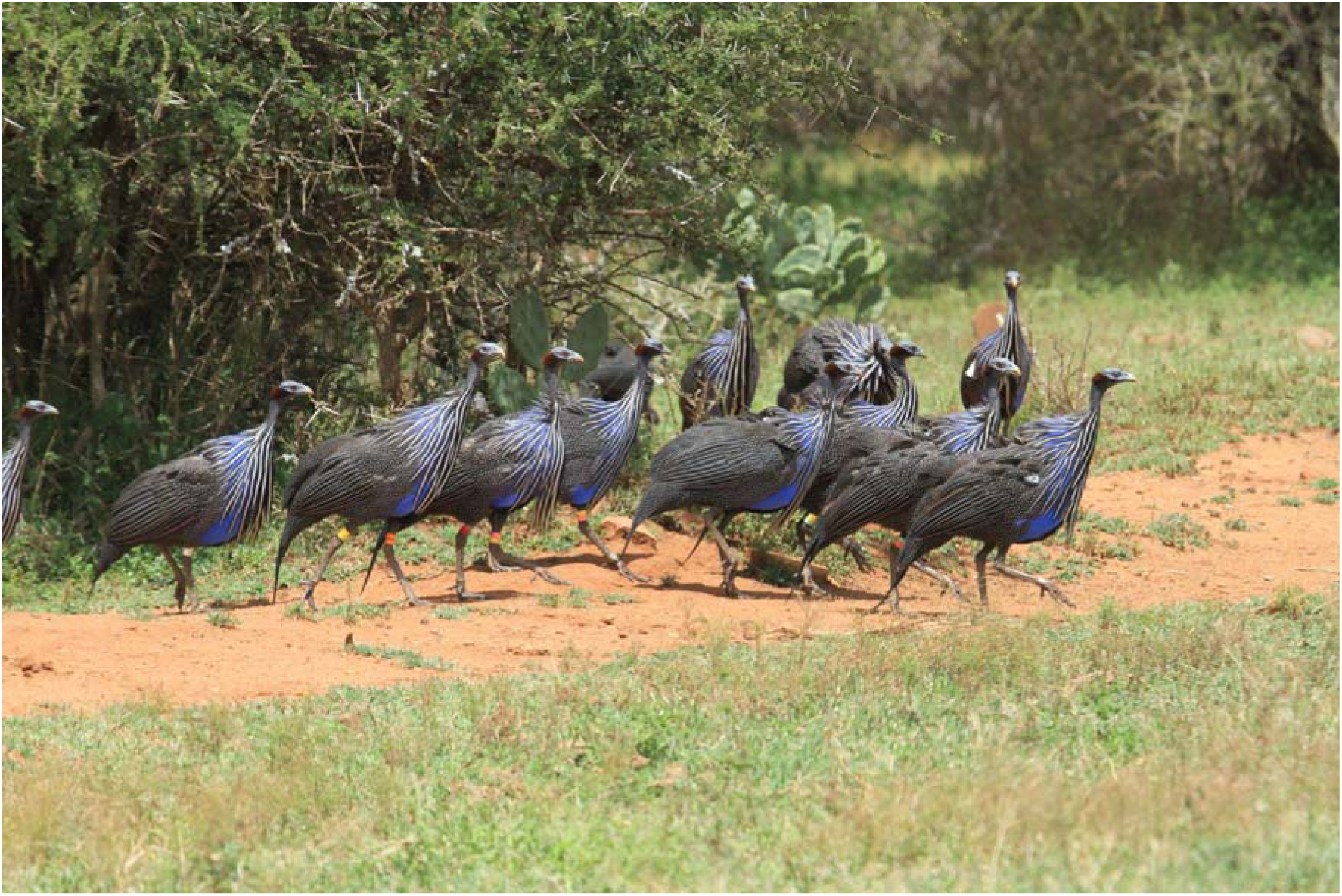
Sub-group of 8 male members from a GPS-tagged group of vulturine guineafowl demonstrating a typical tightly clustered, cohesive movement. Photo: Damien Farine.

## METHODS

### Study system, GPS data collection and pre-processing

We fitted high-resolution solar-powered GPS tags (15 g Bird Solar, e-obs Digital Telemetry, Grünwald, Germany) to all adult members of two social groups of vulturine guineafowl. The first group (Group 1) is composed of 14 males and 11 females, whereas the second group (Group 2) is composed of 7 males and 4 females. GPS tags, together with the harnesses and the platform, which is used to elevate the tag and prevents the solar panel from being covered by feathers, weighed less than 2% of a small bird’s weight, which aligns with biologging ethical guidelines (Portugal & White 2018). GPS tags collected data every fourth day, allowing the battery to charge over the three days that the tags were not operating. The tags were programmed to work from dawn (06:00) to dusk (19:00), collecting high resolution (1Hz) data throughout most of this period. For more details on trapping the birds, tag deployment, programming, and data storage, see Papageorgiou et al. (2019) and for the vulturine guineafowl study system’s design see He et al. (2022). For the present study, we used data collected on six different days between 06/03/2018 and 30/03/2018 for Group 1 and between 16/08/2019 and 05/09/2019 for Group 2, representing a period during which groups maintained a fixed home range and stable group membership.

We excluded one (male) bird of Group 1 and Group 2 from our analyses because of a large number of missing positions. After this removal, we applied linear interpolation to impute missing positions whenever we observe a gap of at most two consecutive fixes for a bird and then extracted all time intervals where we had complete coverage of the positions of all birds, provided these chunks last at least 10 minutes. In total, our dataset comprises 69.1 hours for Group 1 and 17.4 hours for Group 2 (see Supplementary Table 1 for details).

### Calculating dominance ranks

To estimate the dominance hierarchy in the study group, we first recorded all the winners and losers of agonistic associations that we observed, while following the study group on foot for one year [724 agonistic associations collected from January to December 2018, for more details on the association types see Supplementary Table S3 in Papageorgiou & Farine (2020)]. To estimate a robust dominance rank for each individual (Dehnen et al. 2022), we calculated Elo scores (Neumann et al. 2011) by randomising 1000 times the ordering of associations. After creating the 1000 replicated datasets, we calculated the mean and 95% confidence intervals of ranks of individuals. For this process we used the R Package “aniDom” (Sánchez-Tójar et al. 2018; Farine & Sanchez-Tojar 2021).

### Defining associations and instantaneous sub-groups

We defined associations between birds based on GPS proximity. Specifically, for each second, we used GPS coordinates to extract Euclidean distances between all pairwise combinations (dyads) of group members, obtaining a completely connected network. We then filtered the adjacency matrix by dropping all connections whose distance is above a threshold t=5m, and extracted all connected components from the resulting network. Each connected component represents a sub-group of birds that were deemed to be associating with each other at that given time point [equivalent to the gambit of the group (Franks et al. 2010)]. Distance between GPS tags is accurate to within 1m more than 95% of the time (Papageorgiou et al. 2019). We motivate the choice of the distance threshold and provide results for alternative values in the Supplementary Note 1 of the Supplementary Materials. All findings are consistent across the different thresholds.

## Data analysis

### Quantifying the cohesiveness of the base social unit

We assessed how tightly group members stay together. We first identified the largest sub-group at every time and then calculated the percentage of time that this size falls into one of four size classes: 1 to 6, 7 to 12, 13 to 18, and 19 to 24 for Group 1 and 1 to 2, 3 to 5, 6 to 7, and 8 to 9 for Group 2.

Next, we evaluated how individual birds contribute to the overall cohesion of the social unit. For each bird, we measured the fraction of time it was alone – with no other bird within 5m. Then, we built an aggregated association network where edge weights correspond to the simple ratio index, defined as the fraction of time a dyad was observed in proximity (SRI; Hoppitt & Farine 2018). From this weighted network, we computed node strength (i.e., sum of edge weights incident to a node) as a measure of individual gregariousness, and the weighted clustering coefficient as a measure of how cohesive the birds’ neighborhood is (McAssey & Bijma 2015). We report the distribution of the three metrics separately for males and females. To test whether sex differences are greater than expected by chance, we generated 999 realizations of a randomized network where we shuffle sex labels while preserving the network structure. We computed the difference between the male and female means of the quantities of interest for each realization and estimated the empirical p-value as the proportion of such differences that exceed the observed difference.

### Temporal cohesion of sub-groups as a function of sex composition

Differently from the previous section, we now exploit the temporal dimension to measure, for each occurrence of a sub-group, how long it lasted as a measure of temporal cohesion. When observing a particular sub-group – say, four specific birds – we could measure the duration of their association by simply considering the amount of time the occurrence of that set of birds lasted. However, these same four birds might also associate as part of larger sub-groups, and the duration of such intervals wouldn’t be captured automatically. To address this limitation, for each observed sub-group we enumerated all subsets of birds it contains and retained only those subsets that also appear independently as a proper sub-group immediately before or after the larger sub-group they were generated from (see the Supplementary Note 2 of the Supplementary Materials). We considered subsets of size 12 at most and discarded events whose duration is shorter than 10 seconds to focus on meaningful associations. For every subset of birds, we measured the duration of events where the subset stayed continuously together. Then, we show the complementary cumulative distributions of such durations and compute the fraction of events that lasted more than 5 minutes, stratifying for sex composition.

### Transitions between different sub-group sizes

Having analyzed the temporal cohesion of birds, we now aim at studying how birds transition between sub-groups. This analysis complements the previous investigation on “how long associations last” with “where birds go when association events end”. To do that, we estimated the probability a random bird would experience a transition to a sub-group of size *k’*, provided it was in a sub-group of size *k* right before (Iacopini et al. 2024a). Specifically, let *t* denote the time at which a bird experiences a change of sub-group. We recorded the number of times a bird moves from a sub-group of size *k(t)* to a sub-group of size *k(t+1)*. Because *k(t)* birds experience a change of sub-group at time *t*, we divided each count by *k(t)* to avoid overcounting the number of transitions. Note that this measure does not account for birds remaining in the exact same sub-group in order to reflect sub-group changes rather than persistence. The result is a size-to-size transition matrix, which is normalized by rows to obtain transition probabilities P(k(t + 1) | k(t)). We fitted separate transition matrices for male and female birds to compare the way they move between sub-groups.

### Sex differences in spatial positions within sub-groups

Finally, we tested whether male and female birds occupy different spatial positions within sub-groups. Because birds may change sub-groups frequently, we considered birds’ fixes within 30-second windows, sampled 1000 times uniformly at random, and treated the sub-groups identified within each window as fixed for that interval. For each sampled window, we determined sub-groups as the connected components of birds whose average distance is within 5m. Then, after computing the velocity vector of the center of mass of each sub-group, we calculated for each bird their angular position θ (relative to the sub-group’s velocity vector) and surroundedness index. Angular position θ indicates the extent to which a bird is at the front (θ≈0°) or rear (θ≈180°) of a moving sub-group, while the surroundedness index, defined as one minus the sum of the unit vectors from the focal bird towards all other birds in the sub-group, is a robust measure of spatial centrality (He et al. 2022; Christman & Lewis, 2005). We averaged all these quantities over the 30s window. We restricted the analysis to sub-groups that contained at least one male and one female, and comprised at least four birds, because surroundedness is not meaningful in smaller groups.

To test for sex differences in spatial position, we fitted a linear mixed model:

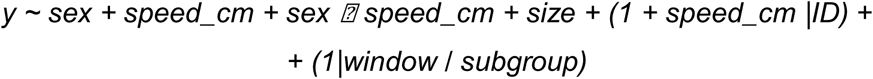

where *sex* refers to the sex of the focal bird (two-level categorical variable, male as reference level), *speed_cm* is the standardized speed of the sub-group’s center of mass, and *size* is the log-transformed size of the sub-group the focal bird is part of. The data have nested (multiple birds per sub-group) and crossed (multiple sub-groups per window) structure. To capture that structure, we included random intercepts and slopes for individual birds *(1 + speed_cm|ID)* and sub-groups within windows *(1|window / subgroup)*. The random effects on individual birds allow modeling birds with heterogeneous associations and response to the sub-group speed. We dropped the latter random effect because it captured negligible variance in the data, supported by a likelihood ratio test (see the Supplementary Note 3 of the Supplementary Materials for details). As a response variable, we used the arctangent of the cosine of θ, to account for the circularity of this metric, and the logit-transformed surroundedness index.

Lastly, we assessed heterogeneities in individual birds’ spatial location within sub-groups by extracting and visualizing individuals’ best linear unbiased predictor (BLUP) from the mixed effect models (Hertel et al. 2020). BLUPs can be considered as indication of among-individual behavioral variation (Hertel et al. 2020; Houslay & Wilson, 2017). We plot the BLUPs corresponding to the random intercept and slope for the two models separately.

## RESULTS

In the main text, we focus on birds belonging to Group 1 because we obtain remarkably consistent results across both groups. We provide detailed results for Group 2 in Supplementary Fig. 1-2.

### The social unit is cohesive with a sex-biased organization

The group spend the vast majority of daylight hours in medium to large sub-groups (Fig. 2a). The largest sub-group consists of more than six birds 89% of the time and is almost evenly split across the three intermediate size strata (7-12 birds: 30%; 13-18: 29%; 19-24: 29%). The group is fragmented into sub-groups of size six or smaller in only 11% of the time. This observation indicates that the group is cohesive and medium to large sub-groups are the typical associative structure (Fig. 3).

**Figure 2.**
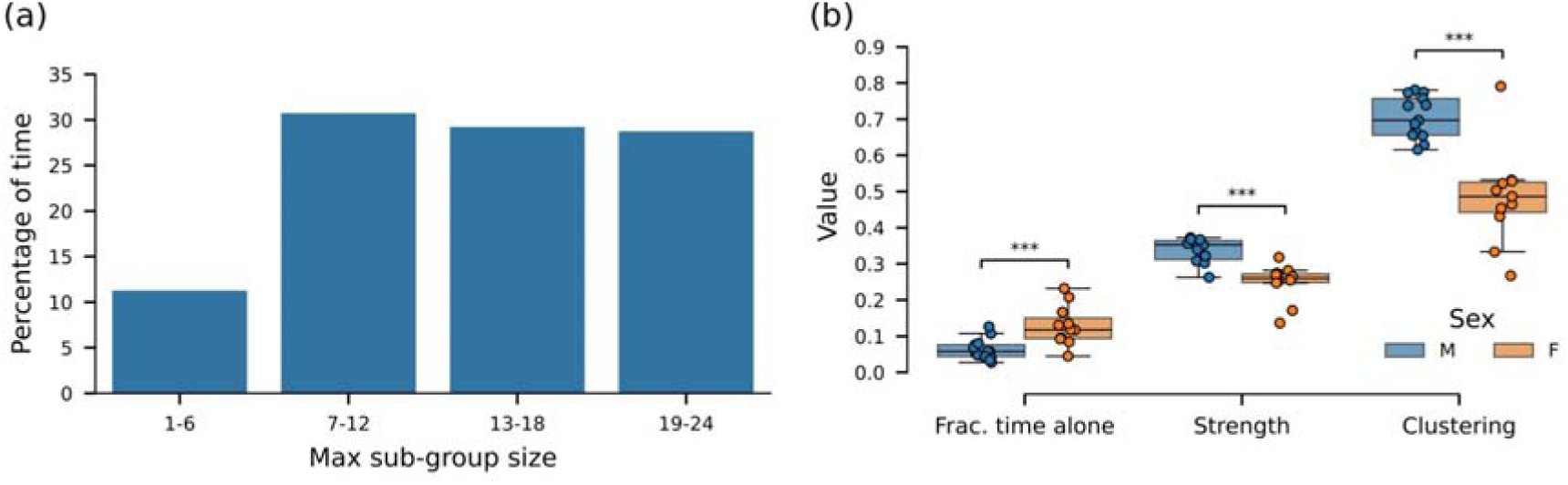
The social group is cohesive, with male and female birds having different positions in the social network built from dyadic relationships. (a) Percentage of time the instantaneous largest sub-group falls into one of four size classes. The largest sub-group is composed of at least seven birds for 89% of the time, indicating strong overall cohesion. (b) Distribution of fraction of time spent alone, node strength, and weighted clustering coefficient, separately for male and female birds. The between-sex difference is significant against permutations of the sex label (stars: *** p < 0.001). Box plots indicate median (middle line), 25th, 75th percentile (box) and 5th and 95th percentile (whiskers), while dots represent individual birds.

**Figure 3.**
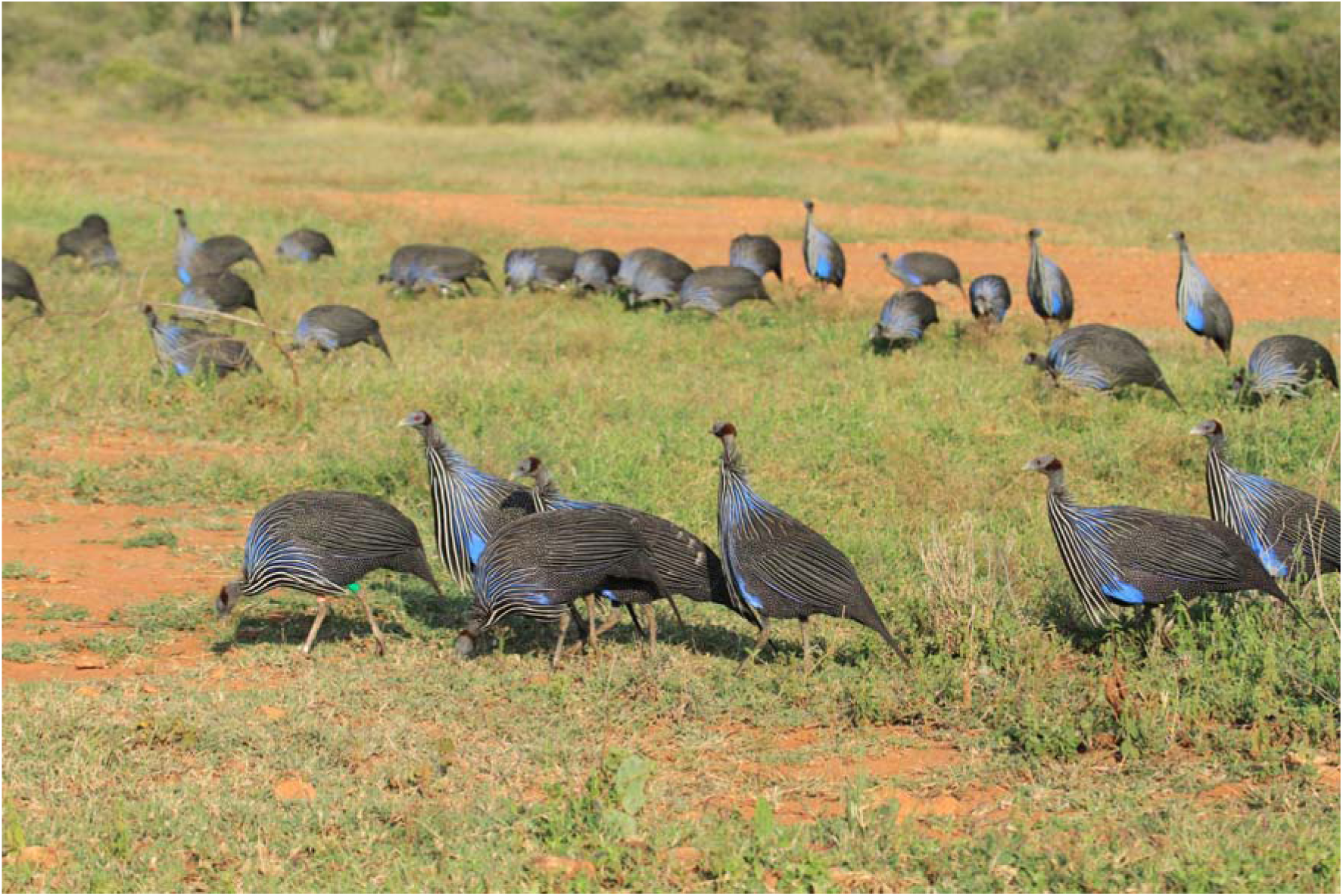
The typical movements of a group of vulturine guineafowl comprises overall cohesion (all group members are within a 20-50 m area) but with distinct clusters of individuals that are extremely close to one-another. The 7 birds in the foreground are all male. Because vulturine guineafowl forage widely distributed resources (e.g. seeds, small insects), spatial patterns do not arise due to differential access to resources. Photo: Damien Farine.

However, the position of birds in the network differs according to sex (Fig. 2b). Female birds spend on average 13% of the time alone, nearly double of the 6% of male birds (p < 0.001). In the aggregate network, males also exhibit higher node strength and clustering (p < 0.001 in both cases), indicating that they not only associate more strongly with other birds but they do so within denser neighborhoods. While the group as a whole remains cohesive, these results point to the internal structure being composed of a strongly cohesive core of males with females occupying more peripheral positions. Interestingly, the female who deviated the most (having more male-like properties) is the same across the three metrics, suggesting that some females may behave more male-alike in terms of their position in the social organization of the group.

These patterns emerge at an aggregate level, remaining robust across different distance thresholds (see Supplementary Fig. 4-5) and consistent with those observed in Group 2 (see Supplementary Fig. 1 a-b). In the following, we exploit the temporal resolution of the data to better understand how such cohesion is achieved through moment-to-moment association patterns.

### All-male sub-groups last longer than other types of sub-groups

Temporal analysis of sub-groups reveals that those composed exclusively of males (e.g. Fig. 1) tend to last longer. The complementary cumulative distribution of event durations show a heavier tail for all-male subsets, compared to the other sex-combinations (Fig. 4a). Interestingly, the presence of just one female is enough to shift the distributions towards shorter durations. The same pattern persists if we separate the different sizes (see Supplementary Fig. 6). More quantitatively, the fraction of association events that last more than 5 minutes is close to double than for other sex combinations (Fig. 4b). For example, considering subsets of size five, 20% of all-male events last more than 5 minutes, whereas this percentage lowers at 13% when only one female is present and to 7-9% for the other sex combinations. The difference between the duration of all-male and other bird sets become smaller as the distance threshold used to define associations increase (see Supplementary Fig. 7-8).

**Figure 4.**
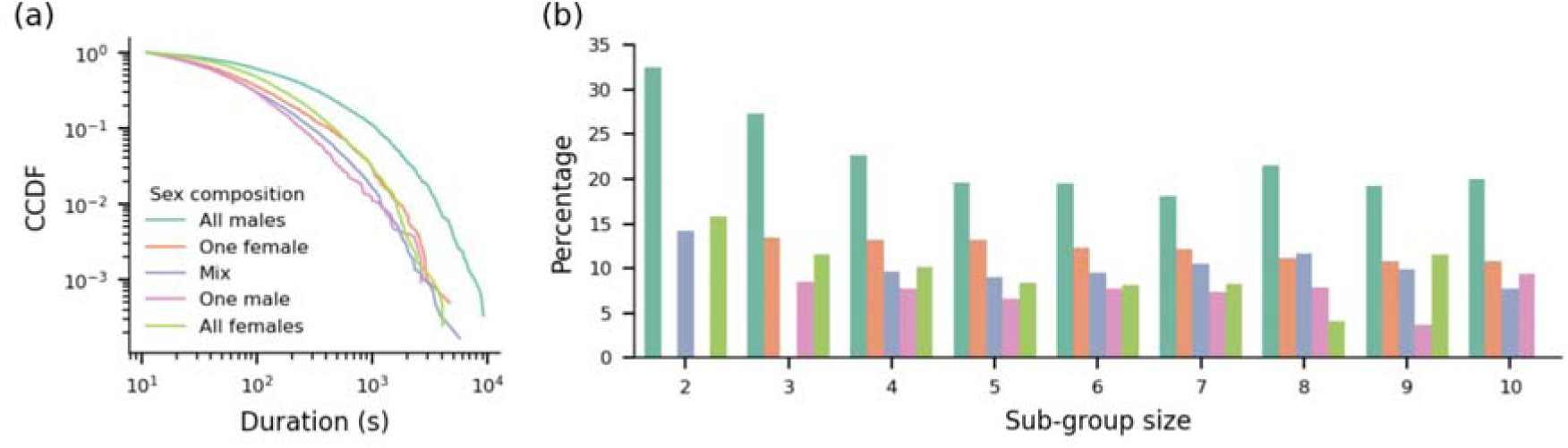
All-male sub-groups last longer than other sex combinations. (a) Complementary cumulative distribution (CCDF) of the duration of association events, separated for different sex compositions. (b) Fraction of bird sets whose duration exceeds 5 minutes, stratified for different sex compositions and sizes.

These observations reveal that the contribution of male birds to the overall group cohesiveness does not merely originate from their aggregate associations, but arises from stable, long-lasting association events. We observe consistent results in Group 2 (see Supplementary Fig. 1 c-d).

### Female birds often transition to isolates

The size-to-size transition matrices highlight sex-based differences in how birds move between sub-groups (Fig. 5). On the one hand, the transition probabilities for male birds are centered around the diagonal (k(t) ≈ k(t+1); Fig. 5a), indicating that the corresponding sub-groups change by losing or gaining a few birds at a time, therefore changing size gradually. On the other hand, female birds not only show a similar concentration of the probability around the diagonal, but also an additional left skewness that peaks in the leftmost column (k(t+1) = 1; Fig. 5b). This means that female birds have a strong tendency to detach alone from sub-groups, regardless of the size – a pattern that remains robust across different choices of the distance threshold (see Supplementary Fig. 9) and is replicated in Group 2 (see Supplementary Fig. 1 e-f). Such solitary departures are likely responsible for the shortened duration of associations involving at least one female bird, and reinforce the picture of male birds being the core of the social unit. These effects are not related to dominance, as the most and least within-sex dominant individuals show the same pattern, with the exception of the male-like female (see Supplementary Fig. 10).

**Figure 5.**
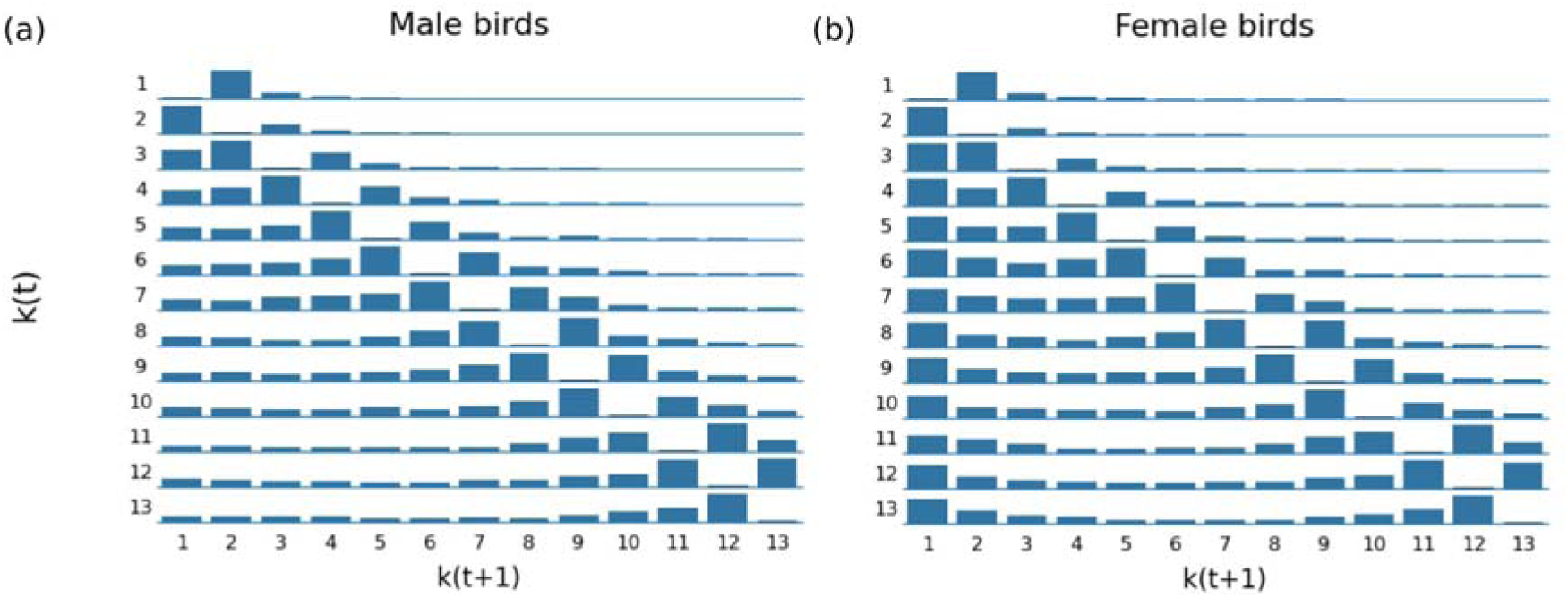
The transition patterns between sub-group sizes are sex-dependent. Each entry of the matrices represents the probability that a bird experiences a transition to a sub-group of size *k(t+1)* conditioned on the bird coming from a sub-group of size *k(t)*. The matrices are fitted separately for male (a) and female birds (b). Female birds show a greater propensity to detach alone from sub-groups, regardless of the size.

### Female birds occupy external spatial positions in sub-groups and move rearwards when in movement

Birds occupy distinct spatial positions within sub-groups. Male birds display larger surroundedness than females (Fig. 6a-c), which indicates that male birds occupy more spatially central positions in sub-groups, whereas female birds stay more external. This picture is similar regardless of the movement speed of the sub-group (Fig. 6a and Fig. 6c). In contrast, front-rear positioning of male and female birds depends on whether the sub-group is in movement or not. Indeed, there is no difference in the distribution of the angular position of male and female birds when the sub-group does not move or moves slowly (speed < 0.1 m/s; Fig. 6b), whereas males tend to the front and females to the rear when the sub-group is in movement (Fig. 6d). These observations are consistent with the linear mixed-model estimates that incorporate additional controls (Fig. 6e). Indeed, both metrics decrease for female birds (fitted coefficients for sex at −0.39 [95% CI: −0.51 – −0.26] and −0.83 [−1.00 – −0.67], respectively), while the angular position of females shift more towards the rear as sub-group speed increases (fitted interaction coefficient at −0.26 [−0.28 – - 0.23]). By contrast, the interaction term for surroundedness was not significant (fitted interaction coefficient at −0.02 [−0.06 – 0.01]). Fitted coefficients are robust against the choice of the distance threshold used to define associations (see Supplementary Fig. 11) and consistent with results from Group 2 (see Supplementary Fig. 1 a-e). Overall, males hold central, frontward positions, whereas females remain spatially peripheral and tend to follow when the sub-group moves.

**Figure 6.**
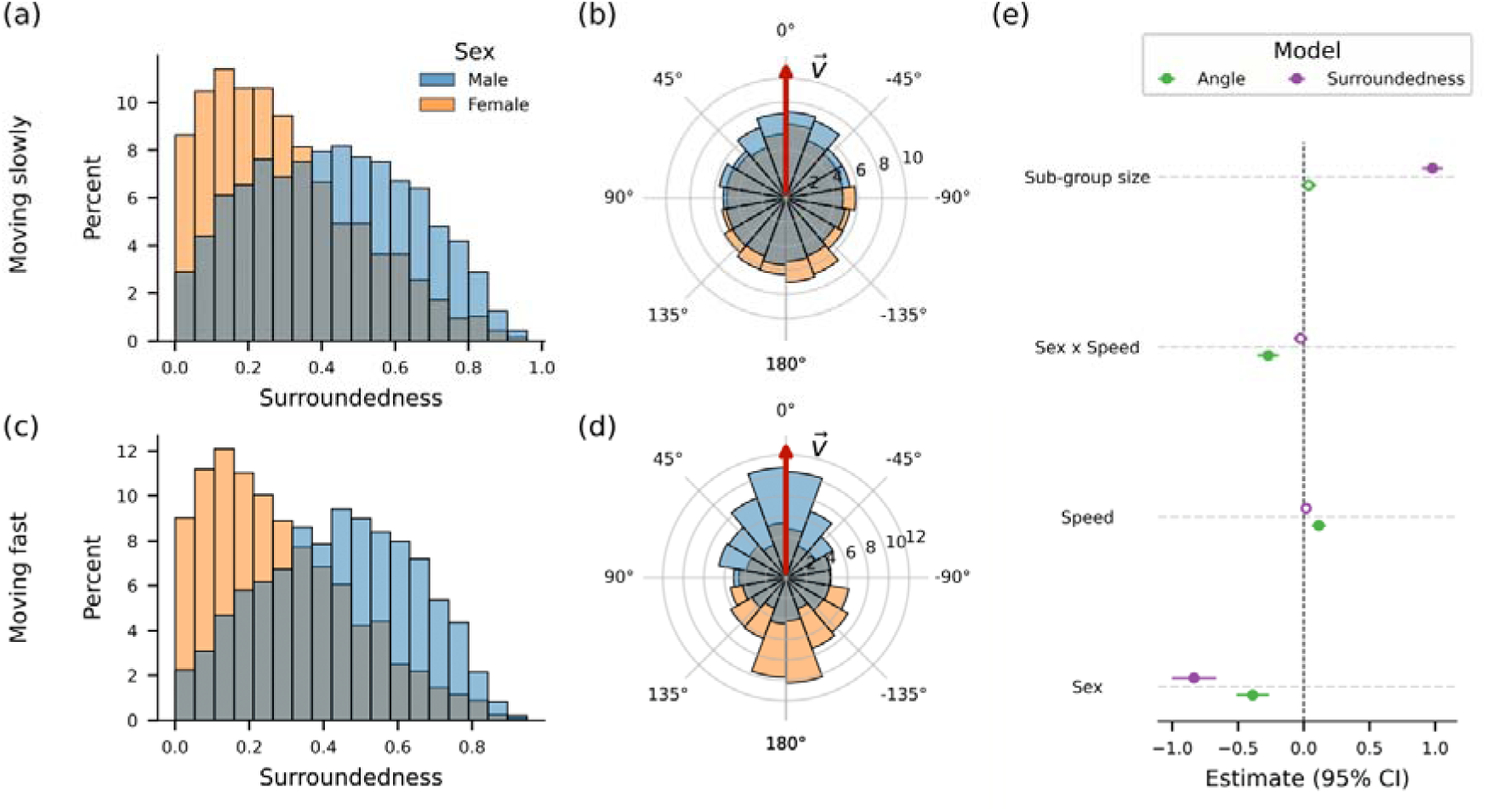
Sex differences in the spatial position of birds within sub-groups. (a-c) Distribution of surroundedness index of male and female birds for slowly moving (a) and fastly moving sub-groups (speed larger than 0.1 m/s; panel c). In both cases, male birds have larger surroundedness, meaning that they occupy more central spatial positions. (b-c) Distribution of birds’ angular position relative to the velocity vector (red arrow), again for slowly moving (b) or fastly moving sub-groups (d). When sub-groups move, male birds occupy the front and females the rear. (e) Summary of the fitted coefficients of the two linear mixed models on the angular position (green) and surroundedness (purple). It shows coefficients’ estimates and their 95% confidence intervals, with full dots indicating significance (p < 0.05). The reference level for sex is ‘Male’. Female birds are associated with a more backward position and lower surroundedness, and they move more towards the rear as the sub-group’s speed increases. Full model results are available in Supplementary Tables 2-3.

Despite the clear sex-dependency in birds’ spatial position, some between-bird differences are present (Fig. 7). For both the angular position and surroundedness models, most birds’ BLUPs are small in magnitude and tightly clustered, indicating that sex already explains most of the variance in birds’ spatial position. Interestingly, one female from Group 1 – the same that was more similar to males in strength and betweenness centrality – is a clear exception. Both its intercept and speed slope BLUPs for the angular position model are larger than the ones of the other females, placing her further frontwards and making her shift rearwards less as speed increases compared to other females. In the surroundedness model, females display a wider variation of intercept compared to males, whereas variation in the speed slope was small for all birds. Again, the same female is among the ones with larger BLUP intercept. We do not observe similar individual deviations in Group 2 (see Supplementary Fig. 2 f-g). Taken together, these results reveal generally low inter-individual heterogeneity with the notable exception of one female from Group 1, whose spatial position within sub-groups is more similar to that of males.

**Figure 7.**
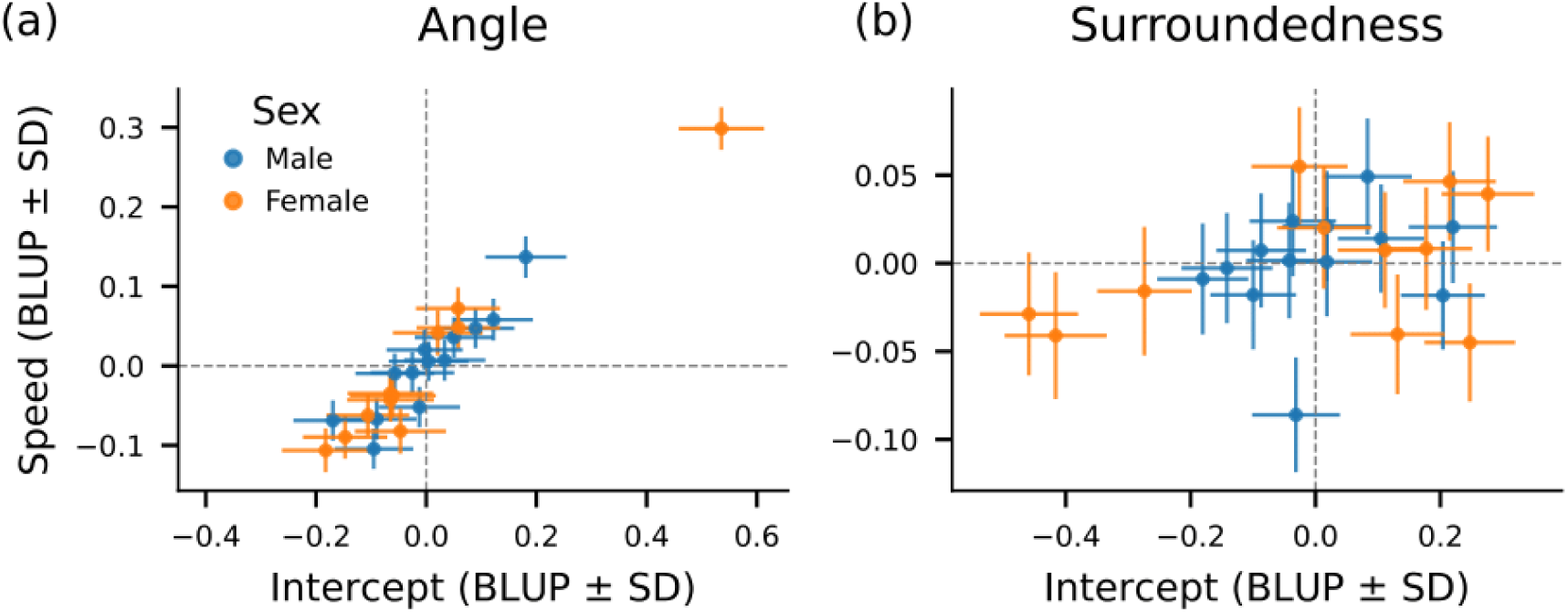
Inter-individual estimates (BLUPs) obtained from the two fitted linear mixed models: (a) regression on angular position and (b) surroundedness. The two axes refer to the birds’ random intercept (x-axis) and random slope (y-axis), and error bars indicate the standard deviation of the estimates. Most of the birds behave coherently within their sex (estimates close to zero). One female makes an exception for the angular position as it deviates towards positive values in both axes, meaning that, on average, it occupies more front position and shifts even more in the front when sub-groups move compared to other female birds. Full BLUP values are available in Supplementary Tables 4-5.

## DISCUSSION

Uncovering the fine-scale structure of cohesive animal groups can be challenging, yet such sub-structure can have major implications for how groups function. Our spatio-temporal modelling of vulturine guineafowl reveals a clear difference among the sexes in their contribution to group cohesion. Combining high-resolution GPS data with higher-order and temporal network approaches, we reveal that males are the central glue of the group by forming tight-knit sub-groups that are temporally stable. By contrast, females orbit more fluidly around this core. These results are consistent across two different groups, thus strengthening the robustness of our findings. Although standard network analyses on the aggregate network already identify sex differences in birds’ centrality, our group-level and temporal approach can capture the underlying dynamics that reveal how these aggregate patterns arise. The patterns that we observe in both groups map directly onto sex-differences in leadership, and our analyses thus reveal mechanisms by which greater influence by males can arise.

Within-group sub-structure is likely to be strongly driven by the long-term relationships among individuals. In social systems where males are philopatric and rarely disperse from their natal group, such as in hamadryas baboons (*Papio hamadryas*) (Jolly 2020) and guinea baboons (*Papio papio*) (Kopp et al. 2015), there should be high genetic relatedness among male group members (Städele et al. 2015). Genetic relatedness, in turn, has long been linked to the strength of social bonds (Hamilton 1964) with kin bonds often setting the core of social systems and enabling cooperation (Städele et al. 2015). In social systems where males are philopatric and set the core of their social group, such as in superb fairy-wrens (*Malurus cyaneus*), males can promote cooperation not only between members of their group but also between groups that used to be united and split after they increased in size (Camerlenghi et al. 2022). In guinea baboons, males that share strong social bonds also support each other when conflicts emerge (Dal Pesco et al. 2022). We found that in groups of vulturine guineafowl, where males are philopatric and thus likely to be highly related to each other (e.g. brothers, father-sons, and cousins), males also form temporally consistent and extremely tight-knit clusters. These clusters form the core of their social group, and thus the glue that holds groups together.

Our analysis of temporal stability of male sub-groups (Figs 4,5) can explain why males were previously found to be more successful at leading the group (Papageorgiou et al. 2024). Vulturine guineafowl, like many other species (including baboons (Strandburg-Peshkin et al. 2015) and fish (Sumpter et al. 2008)), make decisions using a majority rule. Specifically, the outcome of a group decision is determined by the option with the most ‘votes’, or initiators. We found that males are typically found at the front of their sub-groups, which would increase their propensity to be detected as initiators of group movements. However, influence is affected by both initiating *and* being followed (Strandburg-Peshkin et al. 2018). Because male group members are much more consistently found in sub-groups, and these sub-groups are more temporally stable, initiating males will therefore be much more likely to be with other males, placing them in a majority situation. Thus, previously described biases towards male leadership success is unlikely to be because males are inherently more likely to be followed than females, but rather because they maintain stable spatial clusters and thus are more likely to end up as part of a majority. This finding provides an important link between simple dyadic ‘follow-a-friend’ drivers of influence (King et al. 2011) and emergent differences in influence because ‘friends move together’. It has long been suggested that studies of collective decision-making better integrate data on higher-order social groupings when studying influence (Strandburg-Peshkin et al. 2018). Our study confirms this intuition, and provides some of the necessary tools to address this major gap.

Interestingly, we found that one female in Group 1 had higher-order properties that were much more aligned with males than with other females. A plausible explanation here could be that this female had dispersed in the group years ago, bred successfully, and therefore some of the adult males were likely to be her offspring. The pattern that we observed also matches findings that some females from other groups are also highly influential (Papageorgiou & Farine 2020). The fact that only a single female (in this group) expressed this male-like pattern reflects the fact that females can take a number of years to successfully integrate and breed in a group (Nyaguthii et al. 2025; Farine 2025). However, once they successfully breed, they become much more integrated in the group by receiving less aggression from male group members (Dehnen et al. 2025). Therefore, the kin bonds that more dominant females build over time can change their role, eventually holding more male-like social positions. These patterns match the predictions about how within-group relatedness changes with age, specifically increasing during individuals’ reproductive phase of life (He et al. 2025) and is likely to have a number of downstream consequences (e.g. being found more often in the majority is expected to reduce the burden of leadership (Brandl et al. 2025)).

Taken together, our sub-group-level, temporal analysis reveals that the social structure of vulturine guineafowl is predominantly driven by sets of males, and not built-up from distinct and independent dyadic relationships (e.g. male-female mated pairs). Across a broad range of social animals, groups are likely to be composed of more than pairs of individuals that aggregate together, but rather of sets of individuals that together form non-random units. Evidence for this comes from a range of systems, even some in which social associations have long been thought to be expressed relatively randomly. For example, the social network in winter flocks predicts breeding neighbourhoods during the subsequent spring in both great tits (Firth & Sheldon 2016) and blue tits (*Cyanistes caeruleus*) (Beck et al. 2020). In wild Indo-Pacific bottlenose dolphins (*Tursiops aduncus*) that form multi-level alliances, second-order alliances consist of up to 14 individuals including multiple dyads and trios of dolphins (first-order alliances) that move and consort females together. Second-order alliances can also have preferential associations with specific other second-order alliances, forming third-order alliances that can compete against dolphins from rival third-order alliances (King et al. 2021). Finally, bell miners (*Manorina melanophrys*) are socially organised into colonies that are formed from several cooperative-breeding units that also associate preferentially with each other to form coteries (Painter et al. 2000). Such patterns suggest that membership to a group (or a sub-group) represents distinct social units that are not reducible down to dyads, even if the dyads within might still vary in the strength of their social bonds. Thus, sub-group-level temporal analyses provide a powerful way to analyze and interpret cohesive animal social structures.

In broad terms, our sub-group-level approach recovers who associates with whom at a fine temporal resolution and for how long, moving beyond associations measured at the dyadic level only. We demonstrate that addressing social organization through the lens of higher-order networks (Battiston et al. 2020; Iacopini et al. 2024b) can provide important insights into how groups function (in our case how differences in influence between sexes arise). In this context, non-dyadic interactions could be represented by mathematical representations such as hypergraphs, for which a variety of measures and computational tools have recently been developed, from higher-order measures of centrality (Benson 2019) and temporal group persistence (Gallo et al. 2024) to mesoscopic structure such as motifs (Lotito et al. 2022) and communities (Ruggeri et al. 2023). These methods are likely to be increasingly important as advances in biologging (Krause et al. 2013; He et al. 2022) and drone-based (Samad et al. 2025; Koger et al. 2023) methods facilitating the collection of high-resolution and high-precision data that allow these methods to be applied while also generating overly dense networks. Continuing efforts will also be needed to develop methods to analyse animal social networks to follow these advancements (Webber & Vander Wal 2019), including making these methods more spatially explicit (Webber et al. 2023).

## Data and code availability

All data and scripts associated with this manuscript are available on FigShare: https://figshare.com/s/8c78b96868e7dc748341. The repository will be made public upon publication.

## SUPPLEMENTARY MATERIAL

**Supplementary Table 1.**
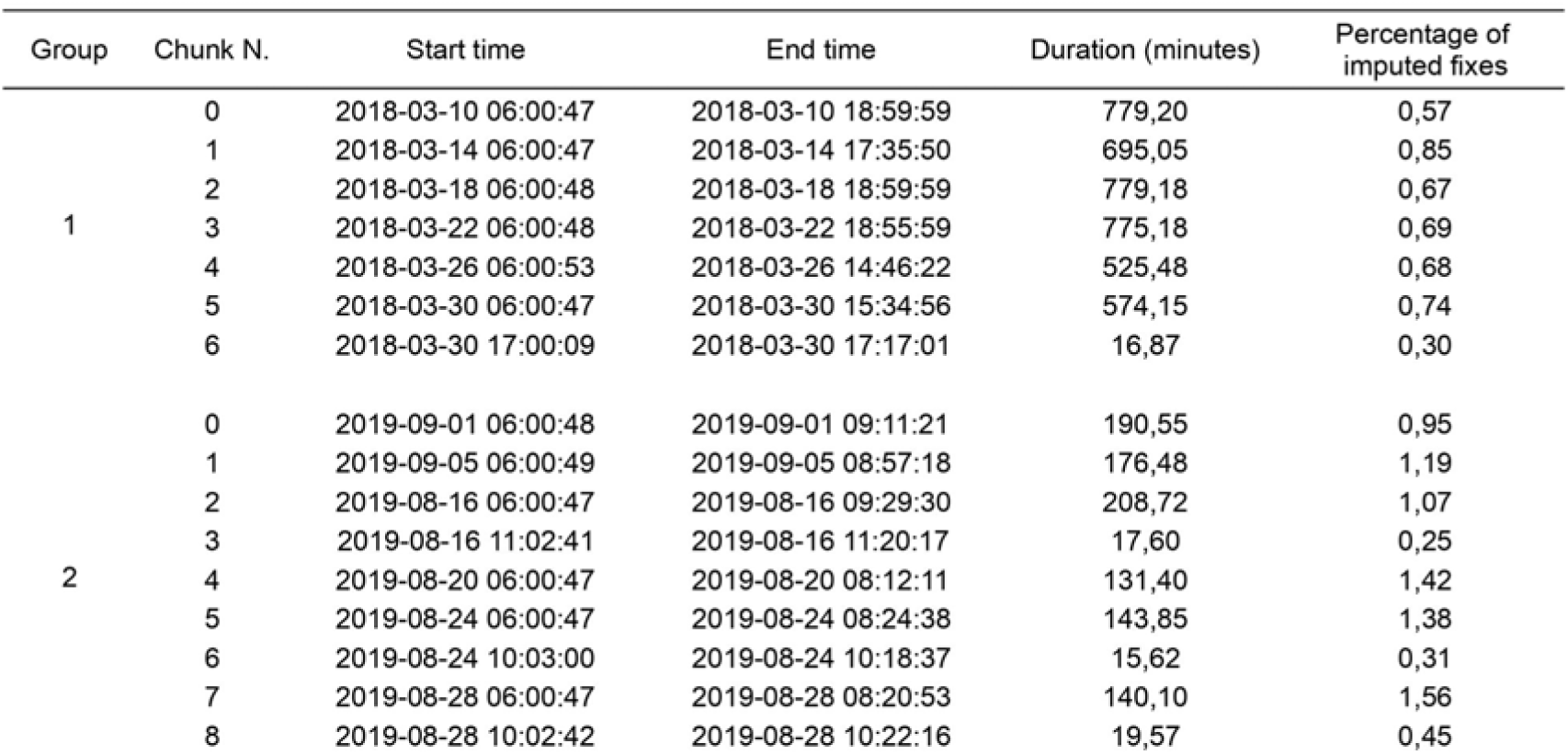
Details on the observation periods used for the analyses. For each group, we extracted all time intervals where we had complete coverage of the positions of all birds. The table shows the start and end time of each of those chunks, with corresponding duration. The last column details the total fraction of GPS fixes that were imputed.

**Supplementary Table 2.** Results of the linear mixed model for the angular position. The reference level of sex is “Male”.

| Angular position |  |  |  |  |  |  |  |
| --- | --- | --- | --- | --- | --- | --- | --- |
| Equation: $\text{arctanh}(\cos(\theta)) \sim \text{Sex} + \text{Speed} + \text{Sex} \times \text{Speed} + \text{Sub-group size} + (1 + \text{Speed} \mid \text{Bird ID})$ | | | | | | | |
| Group 1 |  |  |  |  |  |  |  |
| Fixed Effects Estimates |  |  |  |  |  |  |  |
|  | Estimate | Std. Error | df | t value | Pr(>\|t\|) | 95% CI (lower) | 95% CI (upper) |
| (Intercept) | 0,150 | 0,054 | 55,150 | 2,781 | 0,007 | 0,045 | 0,254 |
| Sex (Female) | -0,388 | 0,063 | 22,250 | -6,130 | < 0.001 | -0,512 | -0,264 |
| Speed | 0,115 | 0,027 | 21,400 | 4,190 | < 0.001 | 0,061 | 0,168 |
| Sub-group size (log) | 0,037 | 0,029 | 16570,000 | 1,289 | 0,198 | -0,019 | 0,095 |
| Sex (Female):Speed | -0,270 | 0,041 | 22,280 | -6,605 | < 0.001 | -0,350 | -0,190 |
| Random Effects Estimates |  |  |  |  |  |  |  |
| Group | Name | Variance | Std.Dev. | Corr. |  |  |  |
| Bird ID | (Intercept) | 0,023 | 0,151 |  |  |  |  |
|  | Speed | 0,009 | 0,094 | 0,93 |  |  |  |
| Residual |  | 0,665 | 0,816 |  |  |  |  |
| Marginal R<sup>2</sup> |  |  |  | Conditional R<sup>2</sup> |  |  |  |
| 0,072 |  |  |  | 0,114 |  |  |  |
| Group 2 |  |  |  |  |  |  |  |
| Fixed Effects Estimates |  |  |  |  |  |  |  |
|  | Estimate | Std. Error | df | t value | Pr(>\|t\|) | 95% CI (lower) | 95% CI (upper) |
| (Intercept) | 0,010 | 0,071 | 626,329 | 0,146 | 0,884 | -0,128 | 0,150 |
| Sex (Female) | -0,341 | 0,034 | 7,271 | -9,968 | < 0.001 | -0,408 | -0,275 |
| Speed | 0,127 | 0,017 | 6,587 | 7,178 | < 0.001 | 0,093 | 0,161 |
| Sub-group size (log) | 0,164 | 0,078 | 5638,578 | 2,096 | 0,036 | 0,010 | 5,17 |
| Sex (Female):Speed | -0,293 | 0,027 | 7,353 | -10,731 | < 0.001 | -0,345 | -0,240 |
| Random Effects Estimates |  |  |  |  |  |  |  |
| Group | Name | Variance | Std.Dev. | Corr. |  |  |  |
| Bird ID | (Intercept) | 0,002 | 0,044 |  |  |  |  |
|  | Speed | 0,001 | 0,031 | 0,290 |  |  |  |
| Residual |  | 0,395 | 0,628 |  |  |  |  |
| Marginal R<sup>2</sup> |  |  |  | Conditional R<sup>2</sup> |  |  |  |
| 0,107 |  |  |  | 0,114 |  |  |  |

**Supplementary Table 3.**
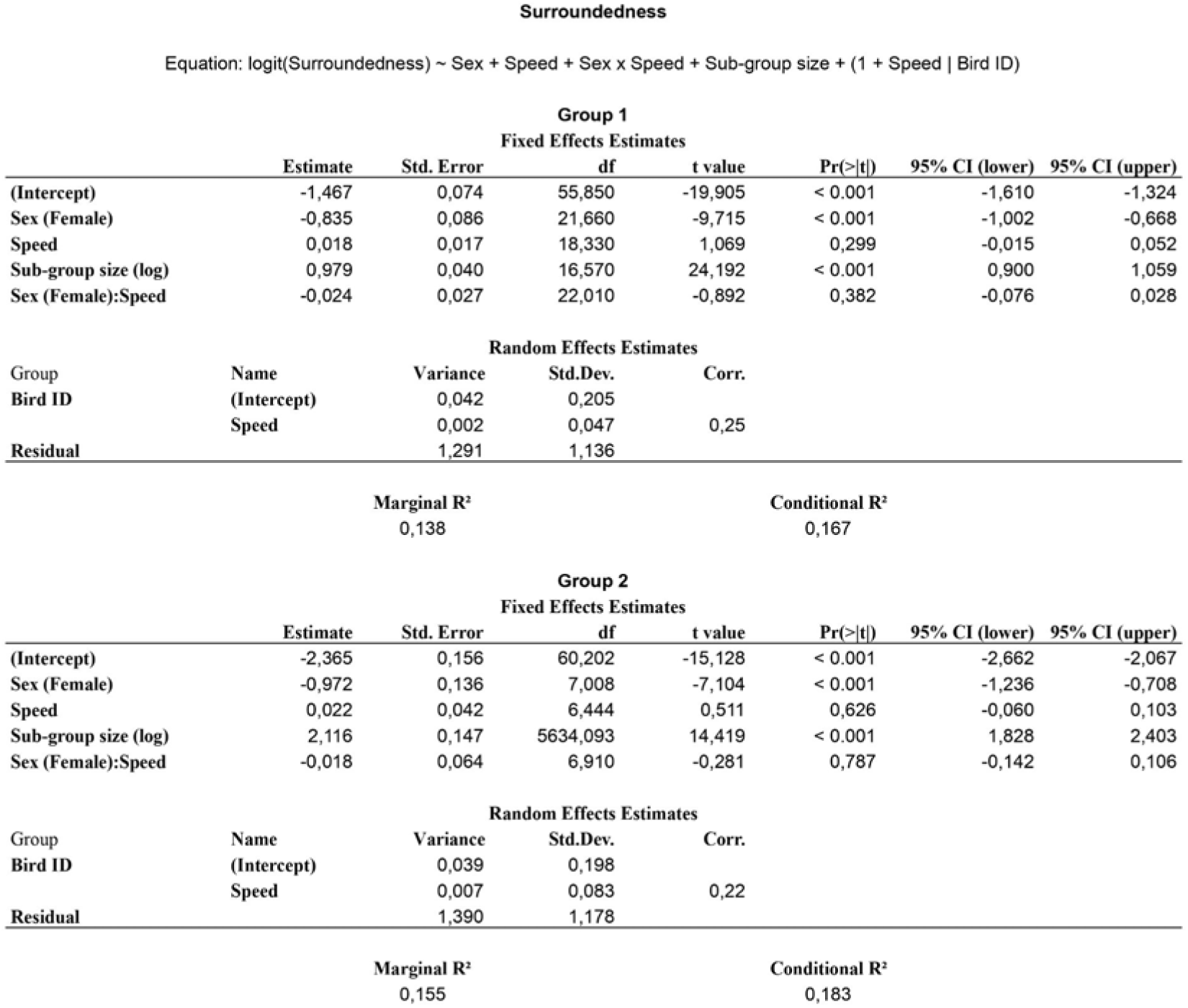
Results of the linear mixed model for the surroundedness. The reference level of sex is “Male”.

**Supplementary Table 4.** BLUP estimates for the individual random effects of the angle model in Group 1. The female that deviates the most from the other females is W1309. We estimated standard deviations (SD) using the REsim function of the merTools package (Knowles & Frederick, 2025).

| Bird ID | Sex | Mean<br>(Intercept) | Median | SD | Mean<br>Speed slope | Median | SD |
| --- | --- | --- | --- | --- | --- | --- | --- |
| W1295 | M | 0,090 | 0,093 | 0,070 | 0,047 | 0,048 | 0,025 |
| W1296 | F | 0,021 | 0,022 | 0,082 | 0,042 | 0,040 | 0,029 |
| W1297 | M | 0,122 | 0,121 | 0,071 | 0,058 | 0,058 | 0,026 |
| W1298 | M | -0,169 | -0,171 | 0,072 | -0,069 | -0,069 | 0,025 |
| W1299 | F | -0,147 | -0,148 | 0,076 | -0,090 | -0,090 | 0,027 |
| W1302 | M | 0,004 | 0,003 | 0,072 | 0,006 | 0,006 | 0,025 |
| W1304 | M | -0,057 | -0,056 | 0,071 | -0,010 | -0,010 | 0,025 |
| W1305 | M | 0,050 | 0,050 | 0,071 | 0,036 | 0,035 | 0,026 |
| W1307 | M | 0,181 | 0,183 | 0,074 | 0,137 | 0,137 | 0,026 |
| W1309 | F | 0,536 | 0,529 | 0,077 | 0,299 | 0,298 | 0,027 |
| W1310 | M | -0,012 | -0,011 | 0,074 | -0,052 | -0,052 | 0,025 |
| W1311 | M | -0,095 | -0,095 | 0,071 | -0,105 | -0,105 | 0,025 |
| W1312 | M | -0,025 | -0,024 | 0,076 | -0,009 | -0,010 | 0,026 |
| W1313 | F | 0,058 | 0,060 | 0,076 | 0,048 | 0,048 | 0,027 |
| W1315 | M | -0,090 | -0,088 | 0,072 | -0,067 | -0,069 | 0,026 |
| W1316 | M | -0,003 | -0,001 | 0,069 | 0,020 | 0,021 | 0,025 |
| W1394 | F | 0,058 | 0,056 | 0,077 | 0,072 | 0,071 | 0,027 |
| W1542 | F | -0,106 | -0,107 | 0,075 | -0,062 | -0,062 | 0,026 |
| W1543 | F | -0,065 | -0,066 | 0,079 | -0,043 | -0,044 | 0,028 |
| W1544 | F | -0,061 | -0,060 | 0,078 | -0,037 | -0,037 | 0,027 |
| W1545 | F | -0,047 | -0,047 | 0,083 | -0,082 | -0,082 | 0,028 |
| W1546 | F | -0,065 | -0,063 | 0,078 | -0,035 | -0,036 | 0,027 |
| W1547 | F | -0,183 | -0,180 | 0,079 | -0,106 | -0,106 | 0,028 |
| W1548 | M | 0,033 | 0,034 | 0,074 | 0,007 | 0,007 | 0,026 |

**Supplementary Table 5.** BLUP estimates for the individual random effects of the surroundedness model in Group 1. We estimated standard deviations (SD) using the REsim function of the merTools package (Knowles & Frederick, 2025).

| Bird ID | Sex | Mean<br>(Intercept) | Median | SD | Mean<br>Speed slope | Median | SD |
| --- | --- | --- | --- | --- | --- | --- | --- |
| W1295 | M | -0,036 | -0,036 | 0,069 | 0,024 | 0,024 | 0,031 |
| W1296 | F | -0,416 | -0,414 | 0,082 | -0,041 | -0,042 | 0,036 |
| W1297 | M | -0,042 | -0,040 | 0,069 | 0,002 | 0,001 | 0,033 |
| W1298 | M | -0,142 | -0,139 | 0,073 | -0,003 | -0,002 | 0,031 |
| W1299 | F | -0,026 | -0,024 | 0,077 | 0,055 | 0,055 | 0,034 |
| W1302 | M | -0,100 | -0,099 | 0,069 | -0,018 | -0,017 | 0,031 |
| W1304 | M | 0,018 | 0,015 | 0,072 | 0,001 | 0,001 | 0,031 |
| W1305 | M | 0,018 | 0,017 | 0,071 | 0,021 | 0,020 | 0,031 |
| W1307 | M | -0,031 | -0,032 | 0,070 | -0,086 | -0,086 | 0,033 |
| W1309 | F | 0,247 | 0,246 | 0,073 | -0,045 | -0,046 | 0,034 |
| W1310 | M | -0,181 | -0,179 | 0,073 | -0,009 | -0,009 | 0,032 |
| W1311 | M | 0,105 | 0,105 | 0,069 | 0,014 | 0,013 | 0,031 |
| W1312 | M | 0,083 | 0,085 | 0,071 | 0,049 | 0,049 | 0,033 |
| W1313 | F | 0,111 | 0,113 | 0,076 | 0,007 | 0,007 | 0,033 |
| W1315 | M | 0,220 | 0,222 | 0,071 | 0,021 | 0,020 | 0,032 |
| W1316 | M | 0,204 | 0,203 | 0,067 | -0,018 | -0,018 | 0,031 |
| W1394 | F | 0,215 | 0,212 | 0,074 | 0,046 | 0,044 | 0,034 |
| W1542 | F | 0,276 | 0,279 | 0,074 | 0,039 | 0,039 | 0,033 |
| W1543 | F | 0,177 | 0,178 | 0,074 | 0,008 | 0,007 | 0,035 |
| W1544 | F | 0,014 | 0,015 | 0,075 | 0,020 | 0,021 | 0,035 |
| W1545 | F | -0,274 | -0,273 | 0,076 | -0,016 | -0,016 | 0,037 |
| W1546 | F | 0,131 | 0,133 | 0,075 | -0,040 | -0,041 | 0,034 |
| W1547 | F | -0,459 | -0,460 | 0,079 | -0,029 | -0,028 | 0,035 |
| W1548 | M | -0,087 | -0,085 | 0,072 | 0,007 | 0,008 | 0,032 |

**Supplementary Table 6.** BLUP estimates for the individual random effects of the angle model in Group 2. We estimated standard deviations (SD) using the REsim function of the merTools package (Knowles & Frederick, 2025).

| Bird ID | Sex | Mean<br>(Intercept) | Median | SD | Mean<br>Speed slope | Median | SD |
| --- | --- | --- | --- | --- | --- | --- | --- |
| W1400 | M | 0,025 | 0,025 | 0,027 | 0,021 | 0,021 | 0,021 |
| W1401 | F | -0,013 | -0,013 | 0,031 | -0,015 | -0,015 | 0,024 |
| W1909 | M | 0,026 | 0,027 | 0,027 | 0,029 | 0,028 | 0,021 |
| W1910 | M | -0,020 | -0,020 | 0,027 | -0,041 | -0,041 | 0,022 |
| W1911 | M | 0,044 | 0,044 | 0,028 | -0,007 | -0,007 | 0,022 |
| W1912 | M | -0,078 | -0,077 | 0,028 | -0,007 | -0,007 | 0,022 |
| W1915 | F | -0,008 | -0,009 | 0,032 | 0,020 | 0,020 | 0,024 |
| W1916 | F | 0,018 | 0,017 | 0,030 | 0,015 | 0,015 | 0,023 |
| W1918 | F | -0,002 | -0,002 | 0,030 | -0,021 | -0,021 | 0,023 |

**Supplementary Table 7.** BLUP estimates for the individual random effects of the surroundedness model in Group 2. We estimated standard deviations (SD) using the REsim function of the merTools package (Knowles & Frederick, 2025).

| Bird ID | Sex | Mean<br>(Intercept) | Median | SD | Mean<br>Speed slope | Median | SD |
| --- | --- | --- | --- | --- | --- | --- | --- |
| W1400 | M | -0,252 | -0,249 | 0,095 | 0,006 | 0,007 | 0,049 |
| W1401 | F | -0,210 | -0,211 | 0,112 | -0,090 | -0,091 | 0,057 |
| W1909 | M | 0,009 | 0,014 | 0,097 | -0,122 | -0,123 | 0,050 |
| W1910 | M | 0,194 | 0,197 | 0,097 | 0,015 | 0,015 | 0,051 |
| W1911 | M | 0,060 | 0,064 | 0,098 | 0,013 | 0,013 | 0,050 |
| W1912 | M | -0,023 | -0,020 | 0,098 | 0,077 | 0,076 | 0,052 |
| W1915 | F | 0,155 | 0,154 | 0,113 | 0,082 | 0,083 | 0,057 |
| W1916 | F | 0,221 | 0,219 | 0,109 | -0,009 | -0,009 | 0,056 |
| W1918 | F | -0,191 | -0,197 | 0,109 | 0,015 | 0,016 | 0,055 |

**Supplementary Figure 1.**
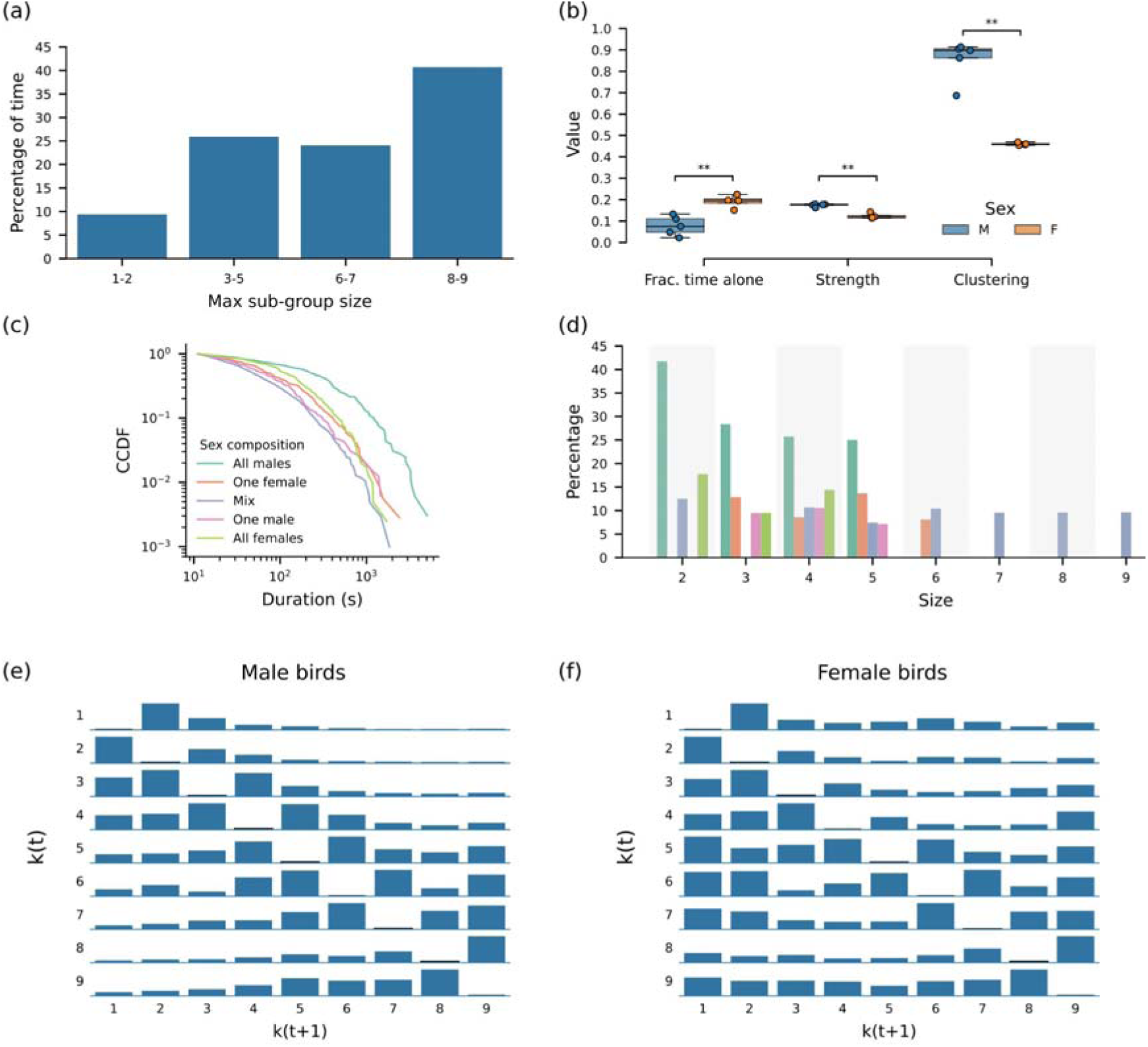
This figure replicates Fig. 2, 4, and 5 of the main text using data of birds in Group 2. (a) Group 2 spent the vast majority of time in medium to large sub-groups, indicating that the group is cohesive. (b) Not all birds contribute the same to the group cohesiveness: male birds spend less time alone and display higher node strength and clustering (p < 0.001). (c-d) All-male sub-groups tend to last longer than sub-groups with any other sex-combination. (e-f) Male birds experience change of sub-groups by losing or gaining a few birds at a time, instead females have a strong tendency to detach alone from sub-groups, regardless of the size of sub-groups they come from.

**Supplementary Figure 2.**
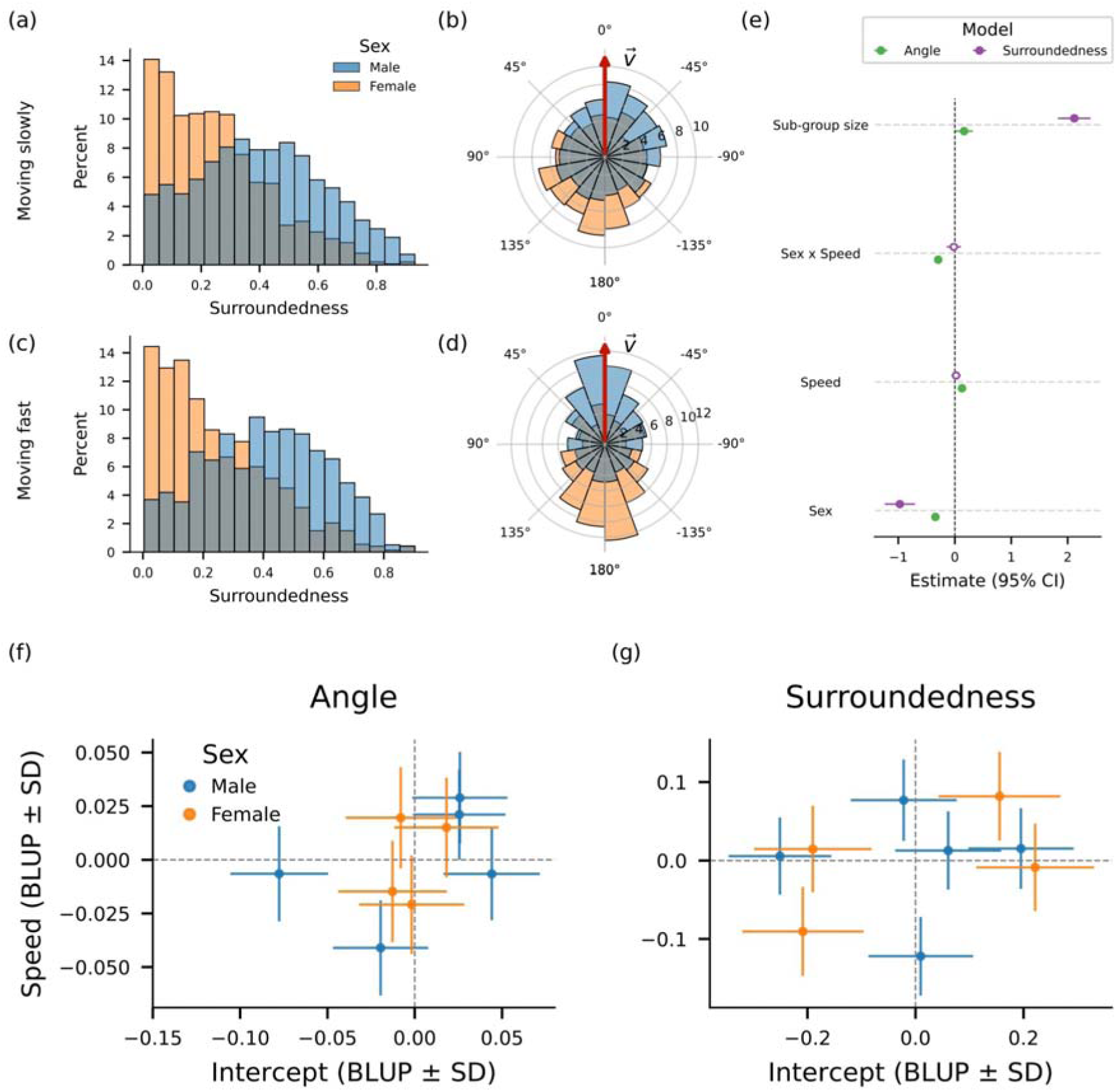
This figure replicates Fig. 6 and 7 of the main text using data of birds in Group 2. (a-d) Male birds occupy more spatially central positions in sub-groups and move to the front when sub-groups are in movement. (e) Results from the linear mixed-model confirm this interpretation while incorporating additional controls. (f-g) We observe no strong heterogeneity between birds. Full BLUP values are available in Supplementary Tables 6-7.

### Supplementary Note 1: Sensitivity to the distance threshold defining associations

Choosing an appropriate distance threshold to effectively capture associations is not trivial. Although many studies indicate that such a choice has to be biologically relevant (Haddadi et al. 2011), there is no consensus on how such a threshold should be determined. Here, we first argue that there is a range of distances that could be considered reasonable and, consequently, we repeat all our analyses for some distance thresholds in that interval. Our findings are robust against the choice of such a distance.

We first have to notice that vulturine guineafowl display a high degree of spatial cohesion, as shown in previous studies (Papageorgiou et al. 2019). The social units under analysis in this study confirm this observation. Indeed, the distribution of pairwise distances peaks below 10m, as shown in Supplementary Fig. 3a,c, although spanning several orders of magnitude, from a few tens of centimeters to around one kilometer for Group 1 and a few hundred meters for Group 2. A complementary picture can be derived from counting the expected number of neighbors a bird has within a given radius, which provides a more macroscopic picture of group cohesiveness. Supplementary Fig. 3b,d shows that the median number of neighbors within a certain radius grows rapidly, with birds having a median of 12 neighbors (i.e., half of the group) already within a distance of 15m for Group 1. For Group 2, birds have a median of 8 neighbors (i.e., all other birds of the group) within a distance of 15m. In addition, the difference between male and female birds is only marginal for distances smaller than 15m.

This description of inter-individual distances suggests that a reasonable upper bound for the distance threshold to capture birds’ associations should be around 10 meters for Group 1, instead for Group 2 that threshold is likely too large. Indeed, above that distance, the social units look like a unique entity with all birds associating together for most of the time, thus leaving small room to study individual preferences across time. Supplementary Fig. 4 shows that, by considering a 10m distance threshold for Group 1, the maximum sub-group size is 24 (i.e., all birds associating together) for more than 20% of the time, and larger than 18 for more than 40% of the time. For the lower bound, we follow Papageorgiou et al. (2024), who set a meaningful distance of 3.5m according to the high degree of spatial cohesion of vulturine guineafowl and to exceed the error of the GPS tags, in which the estimated relative position of two GPS tags is accurate to within 1 m more than 95% of the time (Papageorgiou et al. 2019).

In sum, we repeated our analyses for three distance thresholds within the interval we deem biologically relevant for the species under study, 3.5m, 5m, and 10m, and report in the main text results obtained for the intermediate distance, 5m, for Group 1. Results are consistent across the different distance thresholds as can be seen from Supplementary Figs. 4-11.

**Supplementary Figure 3.**
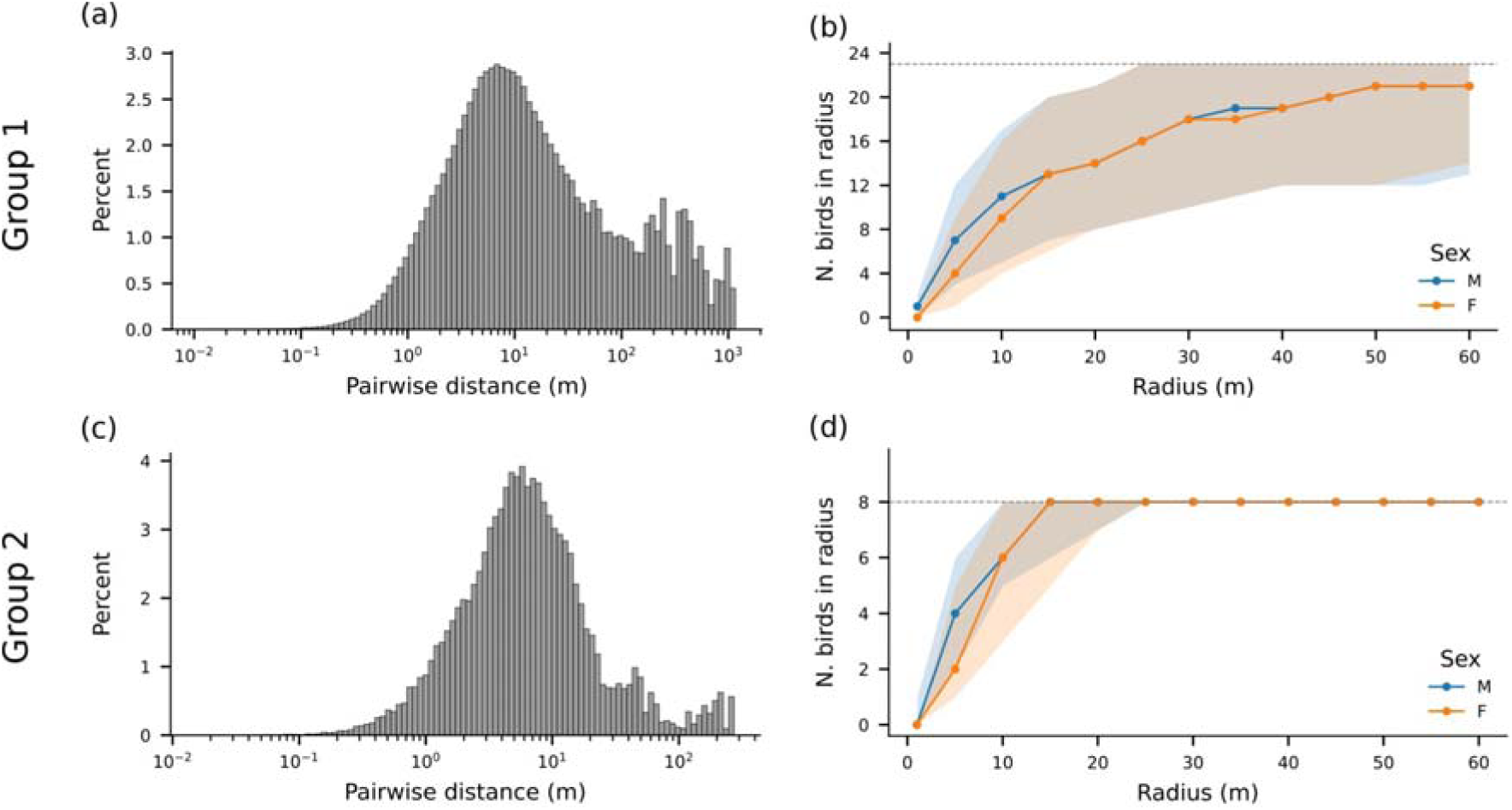
Spatial distribution of the birds within Group 1 (a-b) and Group 2 (c-d). (a,c) Distribution of pairwise distances. (b.d) Number of neighbors a bird is expected to have within a radius (median and inter-quartile range), separated for male and female birds. We sampled birds’ positions every minute. The groups look cohesive. Indeed, birds can expect to find most of the other birds belonging to the group already within 10-20 meters and the distribution of pairwise distances peaks between 1-10m.

**Supplementary Figure 4.**
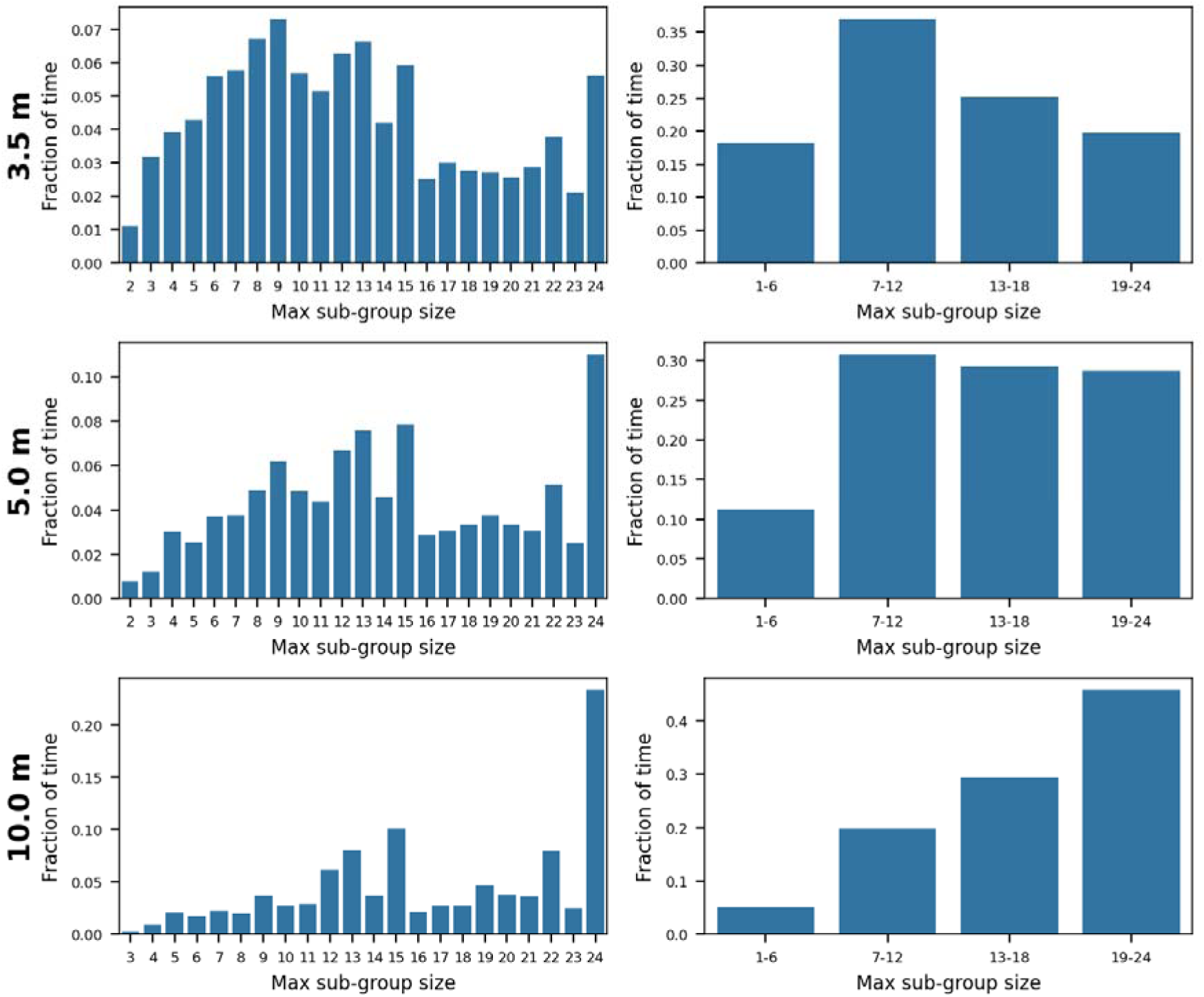
Distribution of time birds in Group 1 are observed in a configuration where the largest sub-group is of a given size, separately for three distance thresholds used to define associations (rows). As the distance threshold increases (from top to bottom), the skewness of the distribution moves towards the right, indicating that already at a 10m distance the social unit starts to look as a unicum (i.e., most of the birds associate together for most of the observation time).

**Supplementary Figure 5.**
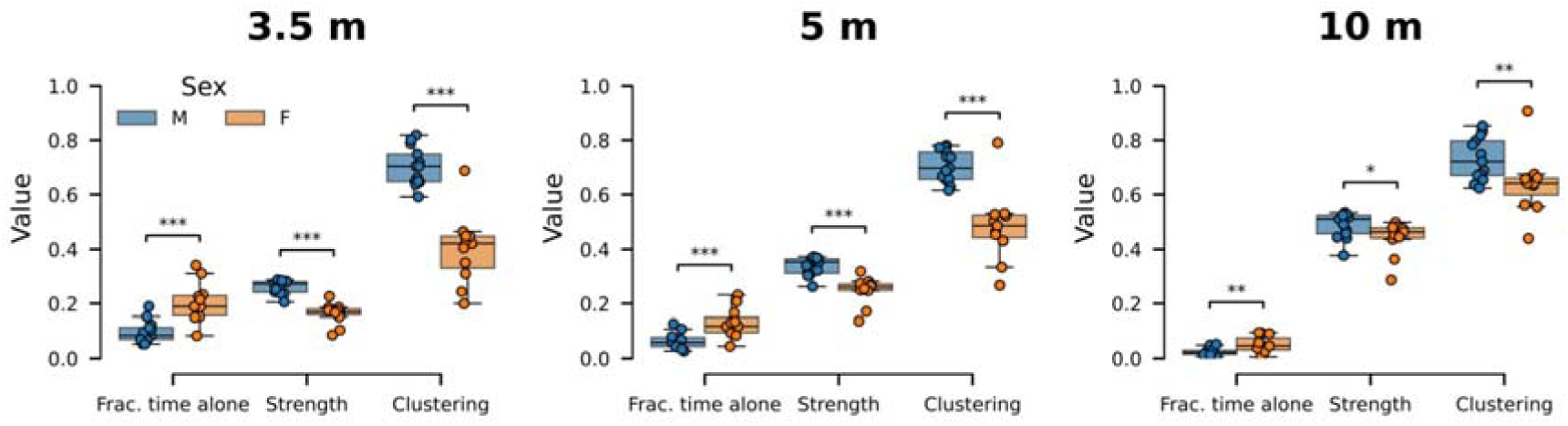
The sex difference observed in Group 1 for the time spent alone, node strength, and weighted clustering coefficient is consistent across the tested distance thresholds defining associations. Stars: * p < 0.10, ** p < 0.05, *** p < 0.001.

**Supplementary Figure 6.**
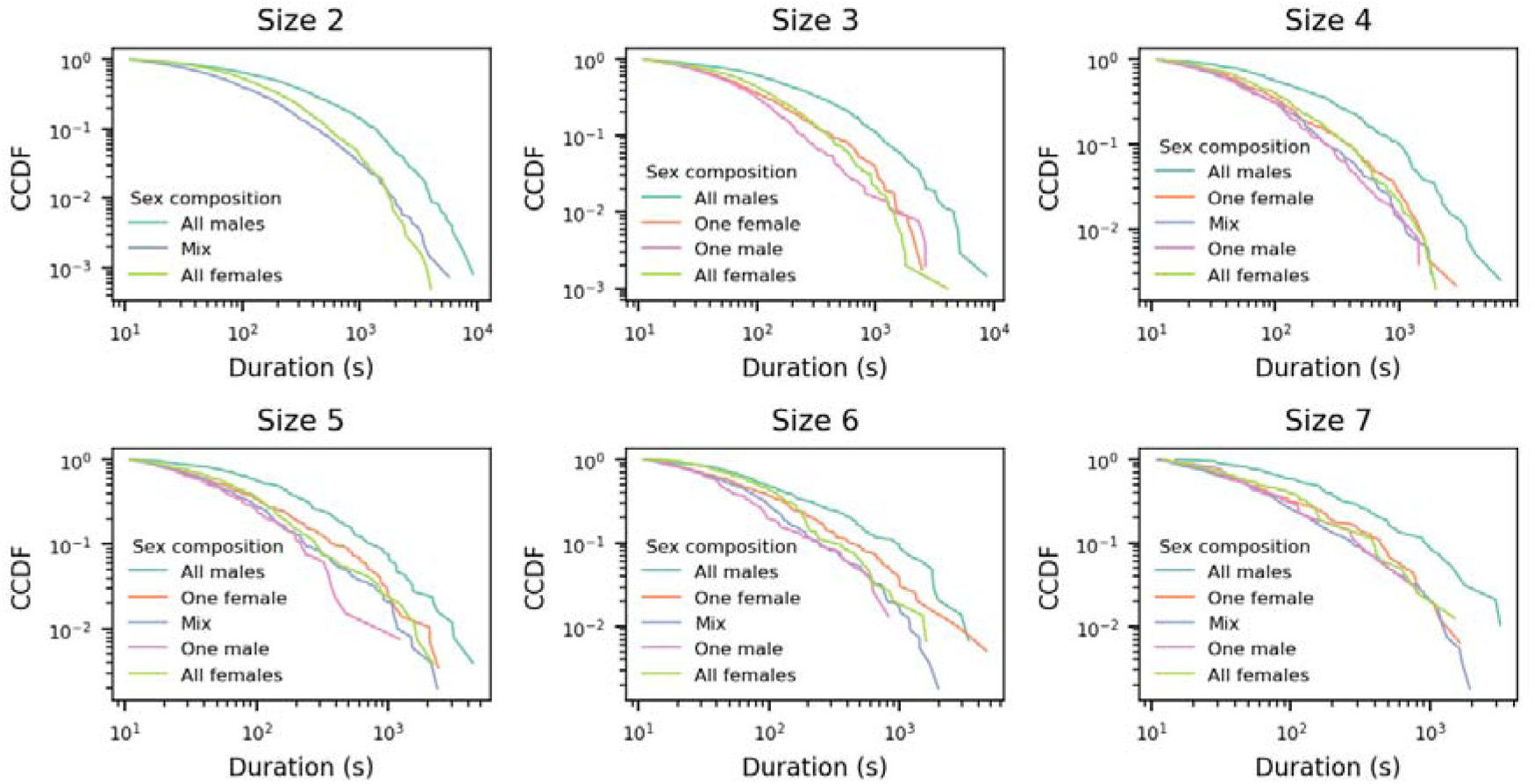
Distribution of bird subset durations observed in Group 1, separated for each subset size. The observation of all-male subsets tend to last longer, as concluded from Figure 2a in the main manuscript, is robust if we look at each subset size separately.

**Supplementary Figure 7.**
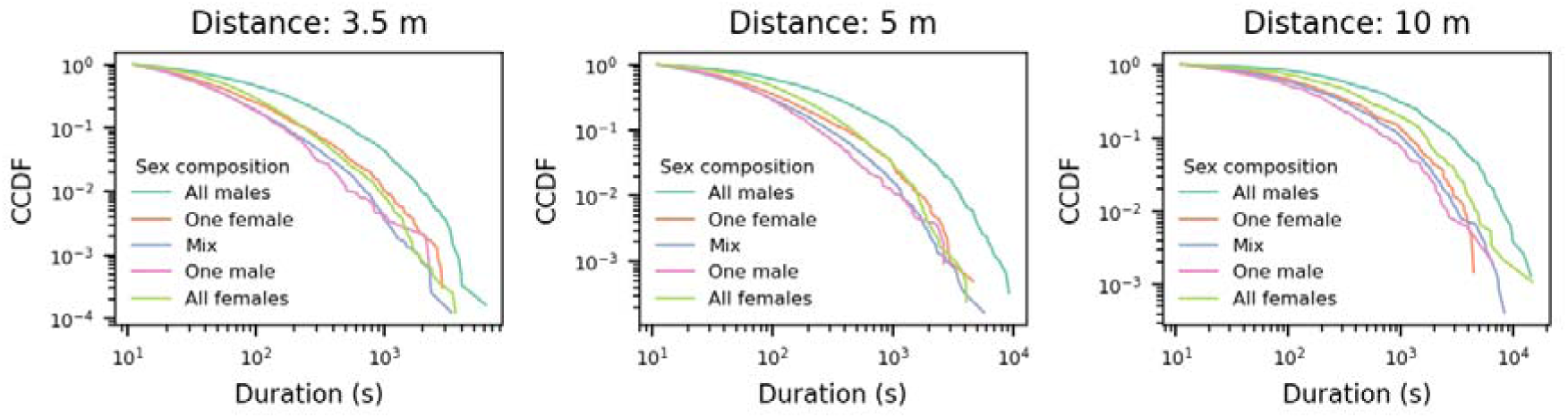
The observation of all-male subsets tend to last longer is robust for different distance thresholds used to determine instantaneous sub-groups in Group 1.

**Supplementary Figure 8.**
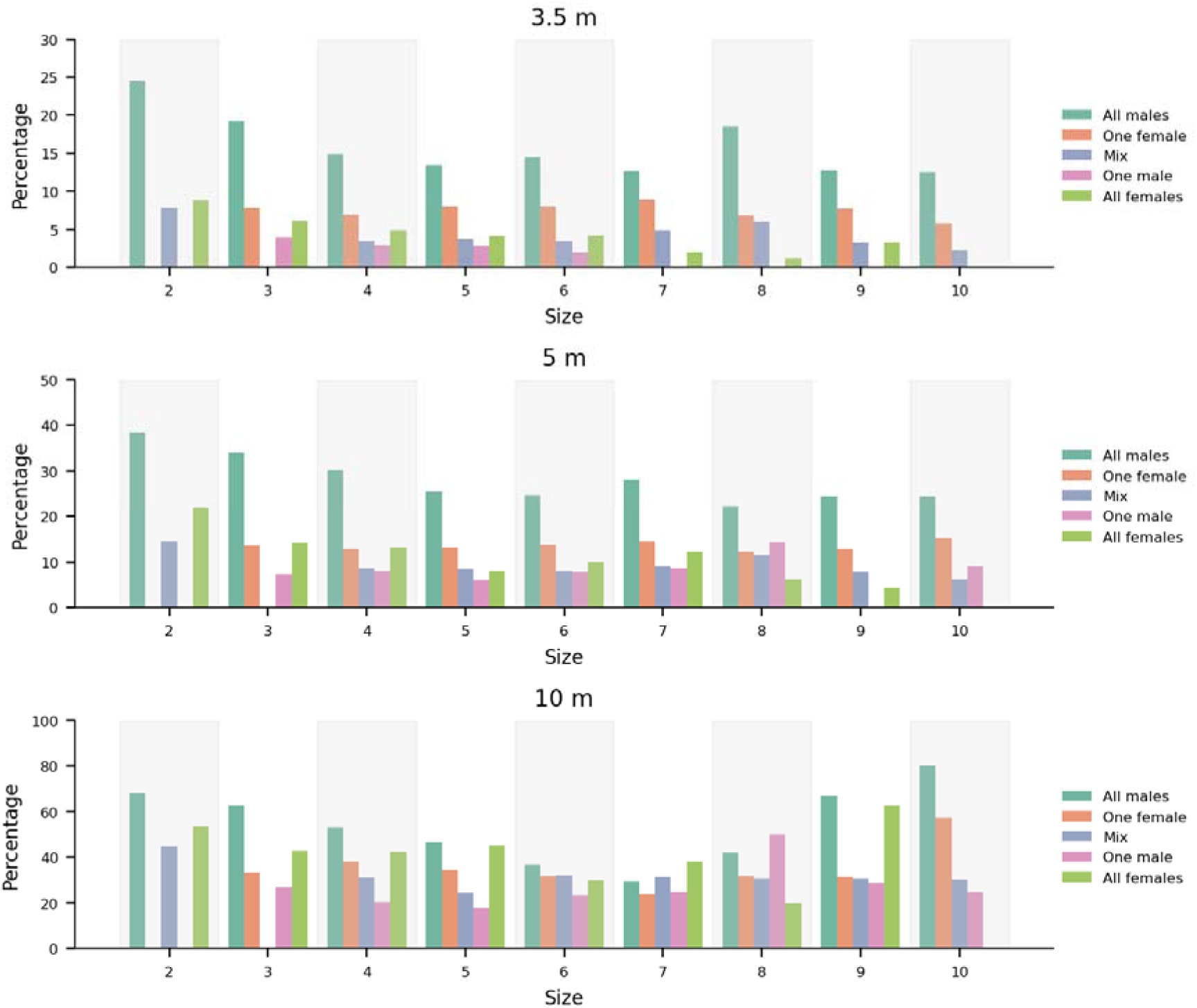
All-male subsets in Group 1 have the largest share of events whose duration exceeds 5 minutes. The observation is consistent across the different distance thresholds employed to define associations. For the largest distance threshold, 10m, all-male subsets do not have the largest share of events above 5 minutes for all sizes.

**Supplementary Figure 9.**
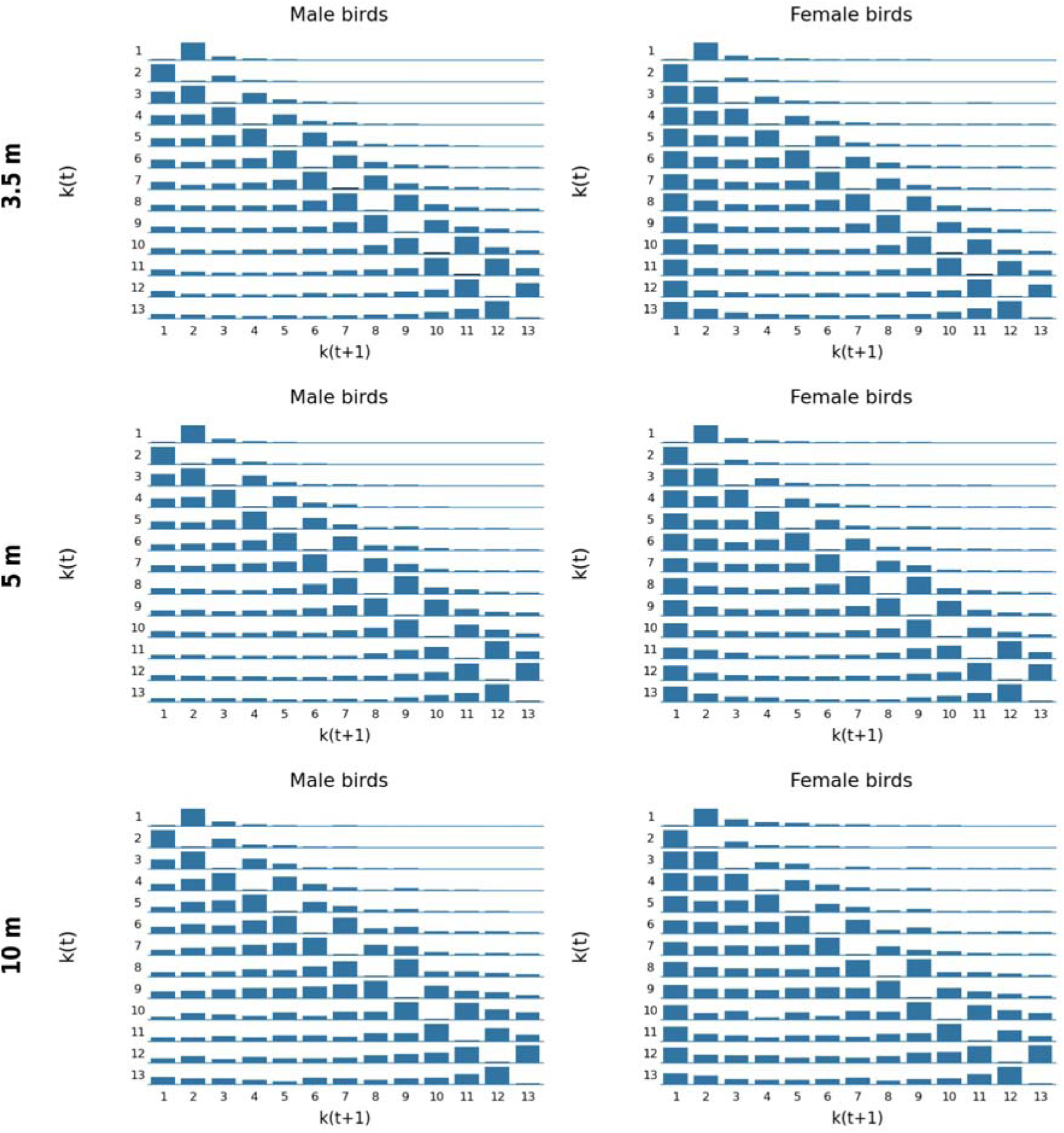
Transition matrices fitted for male and female birds of Group 1 using the different distance thresholds employed to define associations.

**Supplementary Figure 10.**
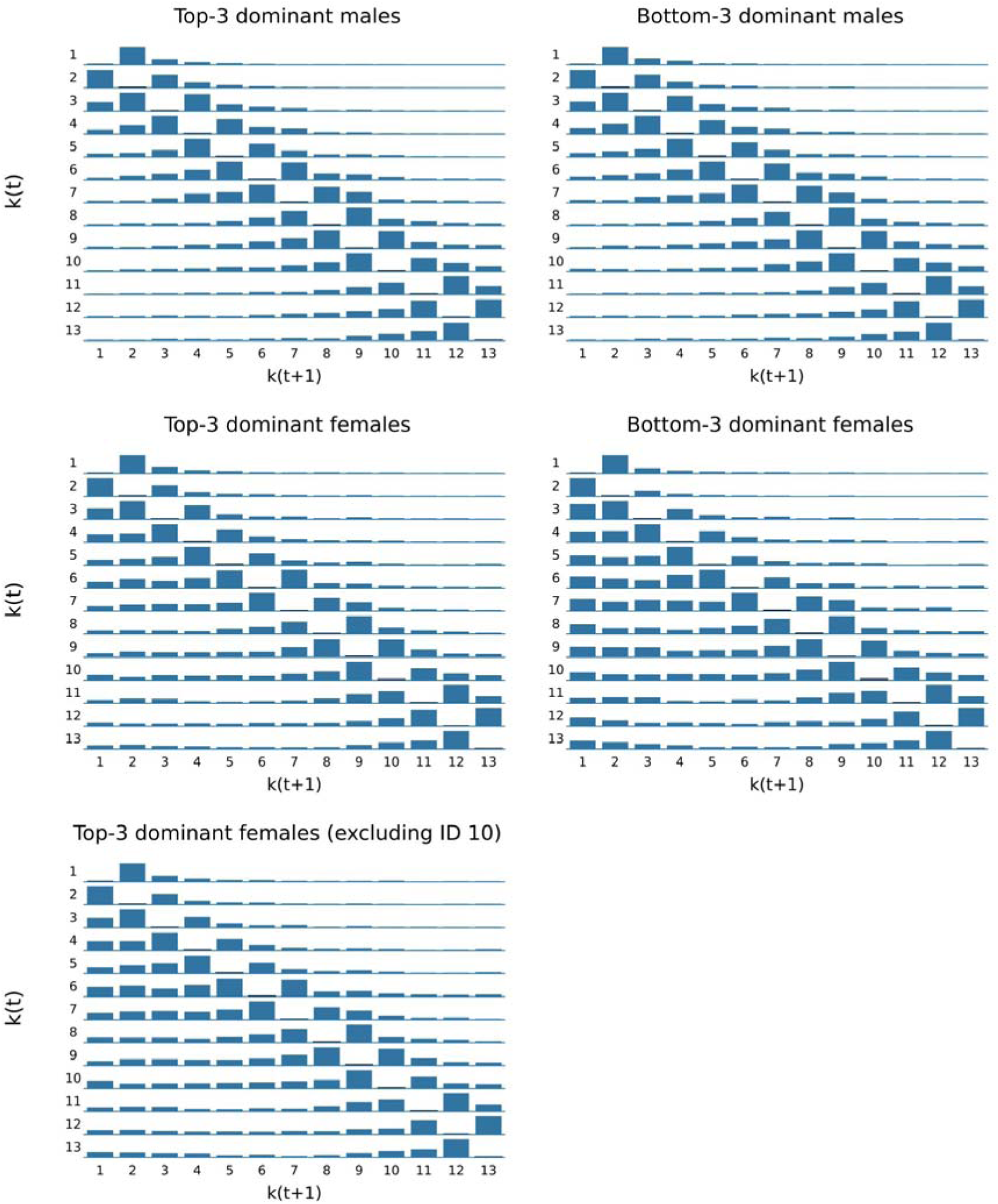
Transition matrices fitted for the top-three (left column) and bottom-three (right column) males and females of Group 1 in the dominance hierarchy. Transition patterns are consistent within sex and distinct between sexes, indicating that the observed sex difference is not explained by dominance rank. The bottom-left panel shows the top-three females after excluding ID 10 (the female with a more front-biased/cohesive pattern).

**Supplementary Figure 11.**
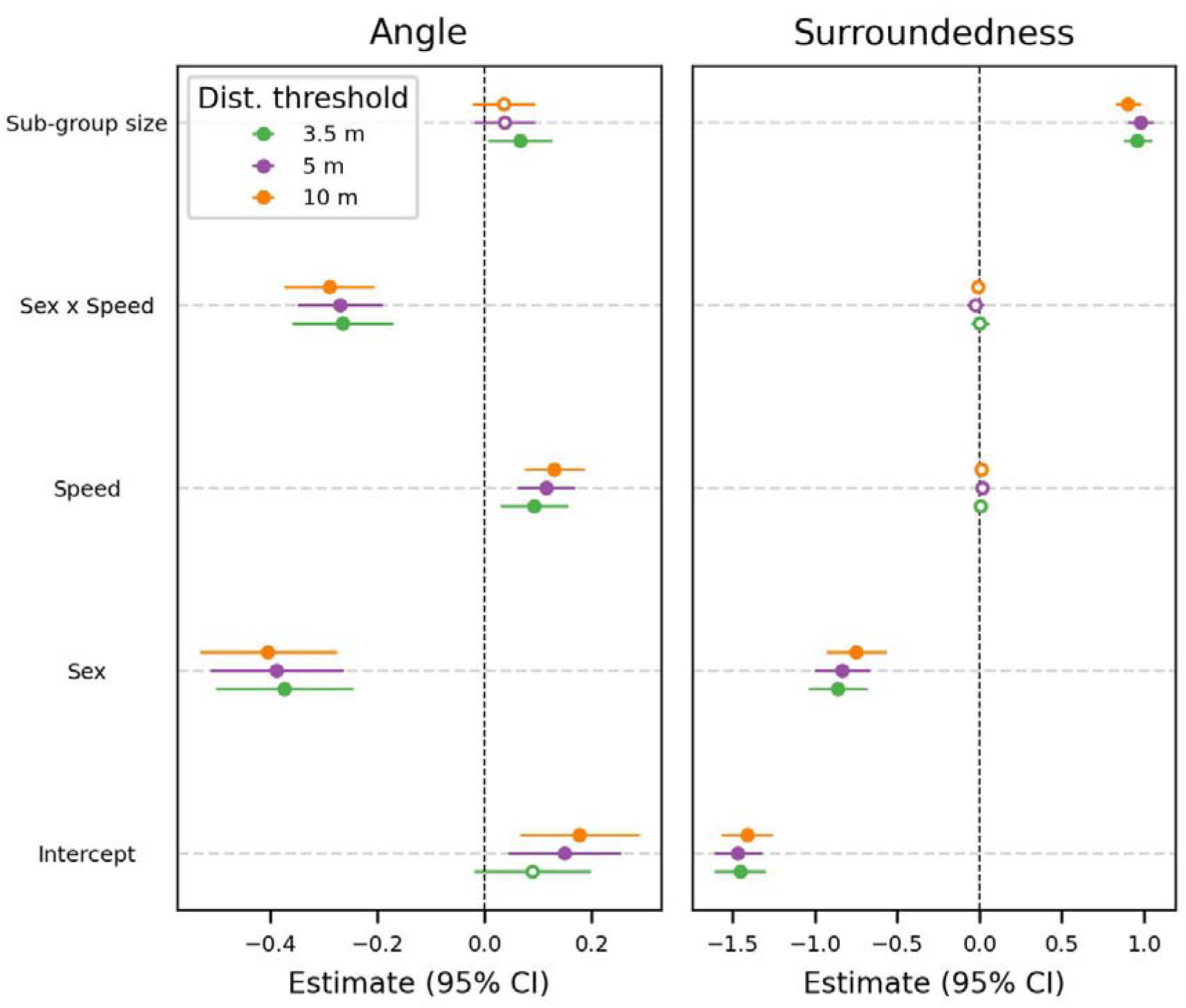
The results obtained from the linear mixed model are consistent across different distance thresholds used to determine sub-groups in Group 1. In the main manuscript, we report the results for the 5m threshold (purple). Note that the surroundedness model for distance threshold at 10m did not converge. The fitted coefficients corresponding to that model were obtained by fitting a model without the random slope. Despite this variation, parameter estimates are consistent with the other models.

### Supplementary Note 2: Details on the method to measure birds’ temporal cohesion

The approach to identify instantaneous sub-groups may fail to effectively track subsets of birds that associate longer because of other birds joining or leaving the sub-group. Take the following sub-group compositions as an example, focusing specifically on birds 1, 2, and 3:

– from t_0_ to t_1_: {(1, 2, 3), (4, 5), (6, 7, 8)}

– from t_1_ to t_2_: {(1, 2, 3, 4, 5), (6, 7, 8)}

– from t_2_ to t_3_: {(1, 2, 3, 4), (5), (6, 7, 8)}

– from t_3_ to t_4_: {(1, 2, 3), (4, 5), (6, 7, 8)}

In this situation, the sub-group (1, 2, 3) is observed only in intervals (t_0_, t_1_) and (t_3_, t_4_), but the triplet of birds actually stays in proximity for the whole observation time (t_0_, t_4_). Thus, by only monitoring the duration of sub-groups, we may overlook sets of birds that stay longer in proximity just because other birds join in the meantime. For this reason, our approach to measure birds’ temporal cohesion does not only consider the instantaneous sub-groups, but accounts for the subsets of birds within each sub-group.

Specifically, the algorithm to measure birds’ temporal cohesion works as follows. For each second, we enumerated all subsets of birds from each sub-group. Then, we retained only those combinations that occur independently of (i.e., not merely as part of) any larger sub-group. In other words, given a subset of birds that originated from a larger sub-group, we look right before and right after the lifetime of the sub-group and check if the subset of birds appeared as a proper sub-group. This logic ensures that we monitor subsets of birds that stayed together not only because they were part of larger gatherings. Considering the previous example, this logic would consider the subset (1, 2, 3) to last from (t_0_, t_4_), whereas the sub-group (1, 2) is not tracked because its occurrence is always within the parent sub-group (1, 2, 3) (i.e., birds 1 and 2 are not observed together in (t_0_, t_4_) if not with bird 3).

The result of this procedure is a list of pairs (t_i, start_, t_i, end_) for each subset of birds, indicating the time interval of the i-th event where the subset is observed in proximity. The corresponding duration of that event is t_i, end_ - t_i, start_. We enumerate subsets of size 12 or smaller, regardless of the size of the parent sub-group. In addition, we then discard all events whose duration is shorter than 10 seconds to focus on meaningful associations.

### Supplementary Note 3: Details on the regression models

As illustrated in the main text, we first selected the random-effects structure using the likelihood ratio test, which supported a model with a random intercept and a random slope of *avg_speed* by individual (ID). Holding this random structure fixed, we then compared a null model (i.e., intercept + random effects only) against the full model, finding the full model provided a significantly better fit according to a likelihood ratio test.

For all models, we performed model diagnostics with DHARMa v0.4.7 (Hartig 2024). Given the temporal structure of the data, we performed two additional checks. First, we checked that the temporal autocorrelation of residuals drops below 0.20 after a few lag values. Second, we fitted the models again by sampling one bird uniformly at random from each sub-group within each snapshot. In such a way, we can get rid of the random factor (1|window / subgroup) because we break the corresponding structure in the data. By repeating this procedure 999 times, we verified that our results are consistent against this robustness test.

